# Real-time visualization of reconstituted transcription reveals RNA polymerase II activation mechanisms at single promoters

**DOI:** 10.1101/2025.01.06.631569

**Authors:** Megan Palacio, Dylan J. Taatjes

## Abstract

RNA polymerase II (RNAPII) is regulated by sequence-specific transcription factors (TFs) and the pre-initiation complex (PIC): TFIIA, TFIIB, TFIID, TFIIE, TFIIF, TFIIH, Mediator. TFs and Mediator contain intrinsically-disordered regions (IDRs) and form phase-separated condensates, but how IDRs control RNAPII function remains poorly understood. Using purified PIC factors, we developed a Real-time In-vitro Fluorescence Transcription assay (RIFT) for second-by-second visualization of RNAPII transcription at hundreds of promoters simultaneously. We show rapid RNAPII activation is IDR-dependent, without condensate formation. For example, the MED1-IDR can functionally replace a native TF, activating RNAPII with similar (not identical) kinetics; however, MED1-IDR squelches transcription as a condensate, but activates as a single-protein. TFs and Mediator cooperatively activate RNAPII bursting and re-initiation and surprisingly, Mediator can drive TF-promoter recruitment, without TF-DNA binding. Collectively, RIFT addressed questions largely intractable with cell-based methods, yielding mechanistic insights about IDRs, condensates, enhancer-promoter communication, and RNAPII bursting that complement live-cell imaging data.

## Introduction

The regulation of human RNA polymerase II (RNAPII) transcription requires an array of proteins and protein complexes, as well as genomic DNA sequences that direct assembly of RNAPII and associated factors to specific genomic loci. The core regulatory assembly for RNAPII transcription initiation is the 4.0MDa pre-initiation complex (PIC), which consists of TFIIA, TFIIB, TFIID, TFIIE, TFIIF, TFIIH, RNAPII, and Mediator.^1^ Note that the PIC includes the TATA-binding protein (TBP), a subunit within the large, 1.3MDa TFIID complex. Over many decades, biochemical experiments have elucidated mechanisms through which the PIC regulates RNAPII initiation.^2–4^ Some basic themes that emerged were that i) transcription *per se* could occur with a minimal system of TBP, TFIIB, TFIIF, and RNAPII;^5,6^ ii) TFIIE and TFIIH worked together to promote RNAPII initiation,^7,8^ in part by melting promoter DNA at the transcription start site (TSS); iii) TFIID helps assemble the entire PIC on promoter DNA, through direct high-affinity contacts downstream and upstream of the TSS;^9^ iv) activated transcription, directed by DNA-binding transcription factors (TFs), requires the 1.4MDa Mediator complex, which triggers TF-dependent RNAPII activation through mechanisms that remain incompletely understood.^10^ The PIC has been structurally characterized using cryo-EM,^11–13^ and structures of actively initiating intermediates have been determined.^14,15^ Consistent with biochemical results,^16,17^ structural data show that TFIID and Mediator cooperate to recruit and orient RNAPII and other PIC factors on promoter DNA.^12^

Over the past 10-20 years, imaging of RNAPII transcription in live cells has transformed our understanding of how gene expression is regulated.^18–20^ Among the many insights provided from live cell imaging are that i) RNAPII transcription occurs in bursts, followed by prolonged dormant periods;^21^ ii) bursts may generate multiple transcripts (i.e. RNAPII initiation and re-initiation from the same promoter);^22,23^ iii) enhancer sequences, which contain many TF binding sites, may activate RNAPII at promoters even without direct enhancer-promoter (E-P) contact;^24–26^ iv) PICs are highly unstable *in vivo*, and Mediator and TFIID appear to be major drivers of PIC assembly and stability.^27^ Finally, v) RNAPII, TFs, Mediator, and other factors form transient clusters in metazoan cell nuclei, which correlate with transcription at nearby genes.^24,28–30^ Such clusters have been reasonably proposed to represent phase-separated condensates,^31^ but verifying condensate formation in cells is challenging and fraught with caveats.^32–34^ Furthermore, directly demonstrating that condensates—in and of themselves—activate RNAPII at specific promoters in cells is not possible with existing techniques.^19,35,36^

With the new mechanistic framework provided by live cell imaging, considered together with foundational biochemical experiments, a series of “next-level” questions have emerged. For example, i) how is RNAPII bursting and re-initiation regulated; ii) how might E-P RNAPII activation occur at a distance (i.e. without direct E-P contact); iii) how do stimulus-response TFs rapidly locate the “correct” target promoters in the nucleus; iv) what are the kinetics of these processes (i, ii, iii), and how are rates impacted by TFs or PIC factors? Finally, v) how do intrinsically disordered regions (IDRs) contribute to RNAPII activation kinetics? Are IDR-containing condensates required for RNAPII activation? Transcription regulatory proteins are enriched in IDRs,^37^ including especially TFs and Mediator.^38,39^ Is their primary function to drive condensate formation, or do they act independently of this biophysical phenomenon?

To address these and other mechanistic questions, we developed a real-time *in vitro* fluorescence transcription assay, which we call RIFT. Key features of this experimental system include i) reconstitution of regulated, TF-dependent transcription with purified human PIC factors (no extracts) at the native HSPA1B promoter, ii) direct visualization and quantitation of nascent RNA transcripts in real time, iii) direct visualization of individual promoters, TFs, or condensates with single-molecule resolution, iv) precise timing, with onset of transcription occurring upon addition of NTPs, and v) continuous monitoring of transcriptional output at hundreds of individual promoters over time, to allow accurate assessment of RNAPII initiation and re-initiation events.

The RIFT assay builds upon pioneering work from the Tjian lab that showed it was feasible to reconstitute and visualize RNAPII transcription at individual promoters. In this case, PICs were assembled with purified factors on promoters immobilized on microscopy slides. After NTP addition, transcripts were visualized by subsequent incubation with fluorophore-tagged DNA sequences that hybridized with RNA products.^40^ More recently, the Buratowski and Gelles labs pioneered methods to visualize PIC factor recruitment to single promoters in real time, using yeast nuclear extracts from cells expressing labeled proteins.^41,42^ This co-localization single-molecule spectroscopy (CoSMoS) system has markedly advanced understanding about factor recruitment and PIC assembly,^43^ although transcriptional output is not measured with CoSMoS.

We applied the RIFT system toward the “next level” questions cited above. We started with a “standard” set of RIFT experiments followed by assays that mimic cellular “stimulus response” conditions requiring rapid TF-dependent transcriptional changes. We focused on Mediator and TFs (HSF1 in particular) because each was shown to have the greatest impact on RNAPII activity. Our results are consistent with mechanistic models inferred from live cell imaging and single-cell nascent transcriptomics. For example, our RIFT experiments yielded insights about how burst size and duration is regulated, how TFs can rapidly find their target promoters, and how enhancers might activate nearby promoters without direct E-P contact. Furthermore, we establish that all TF- and Mediator-dependent effects on RNAPII activation do not require condensate formation; however, condensates could further enhance RNAPII activation kinetics and/or transcriptional output, and this is an area for future study.

## Results

### Development of a Real-time *In vitro* Fluorescent Transcription (RIFT) assay

The RNA imaging technology utilized for RIFT is based on the Peppers RNA aptamer,^44^ which folds into a secondary structure that binds to fluorogenic small molecules with high affinity. We hypothesized that the Peppers aptamer could be adapted for real-time detection of nascent transcripts as they emerged from an actively transcribing RNA polymerase. As an initial proof-of-concept, we completed *in vitro* transcription with T7 RNAP and used a plate reader to measure bulk fluorescence. These experiments confirmed that fluorescent output was dependent on the fluorogenic ligand, DNA template, and T7 RNAP (**Figure S1A**), demonstrating that the Pepper RNA aptamer could allow *in situ* visualization and quantitation of RNAP transcription.

We next evaluated the Pepper RNA aptamer in our well-tested human RNAPII transcription system, which uses purified human PIC factors (no extracts; **Figure S1B**).^45,46^ To adapt for RIFT, we inserted tandem Peppers aptamer sequences 100bp downstream of the transcription start site on the native human HSPA1B promoter (**Figure 1A**). Thus, all native regulatory sequences (e.g., TATA box, initiator element) were retained. The HSPA1B promoter template was also tethered to biotin at its 5’-end. The regulatory TF for HSPA1B, HSF1, was pre-bound to the template followed by PIC assembly. The biotinylated templates were then added to streptavidin-coated flow-channel microscopy slides, and ribose nucleotide triphosphates (NTPs) were added to initiate transcription (**Figure 1B**); the moment of NTP addition was designated t=0. To visualize RNAPII transcription in real time, continuous single-molecule Total Internal Reflection Fluorescence (smTIRF) imaging was conducted out to 3 minutes (**Figure 1C**), with a frame rate of 5/sec (200msec; 400msec for 2-color imaging) for precise temporal resolution.

**Figure 1.**
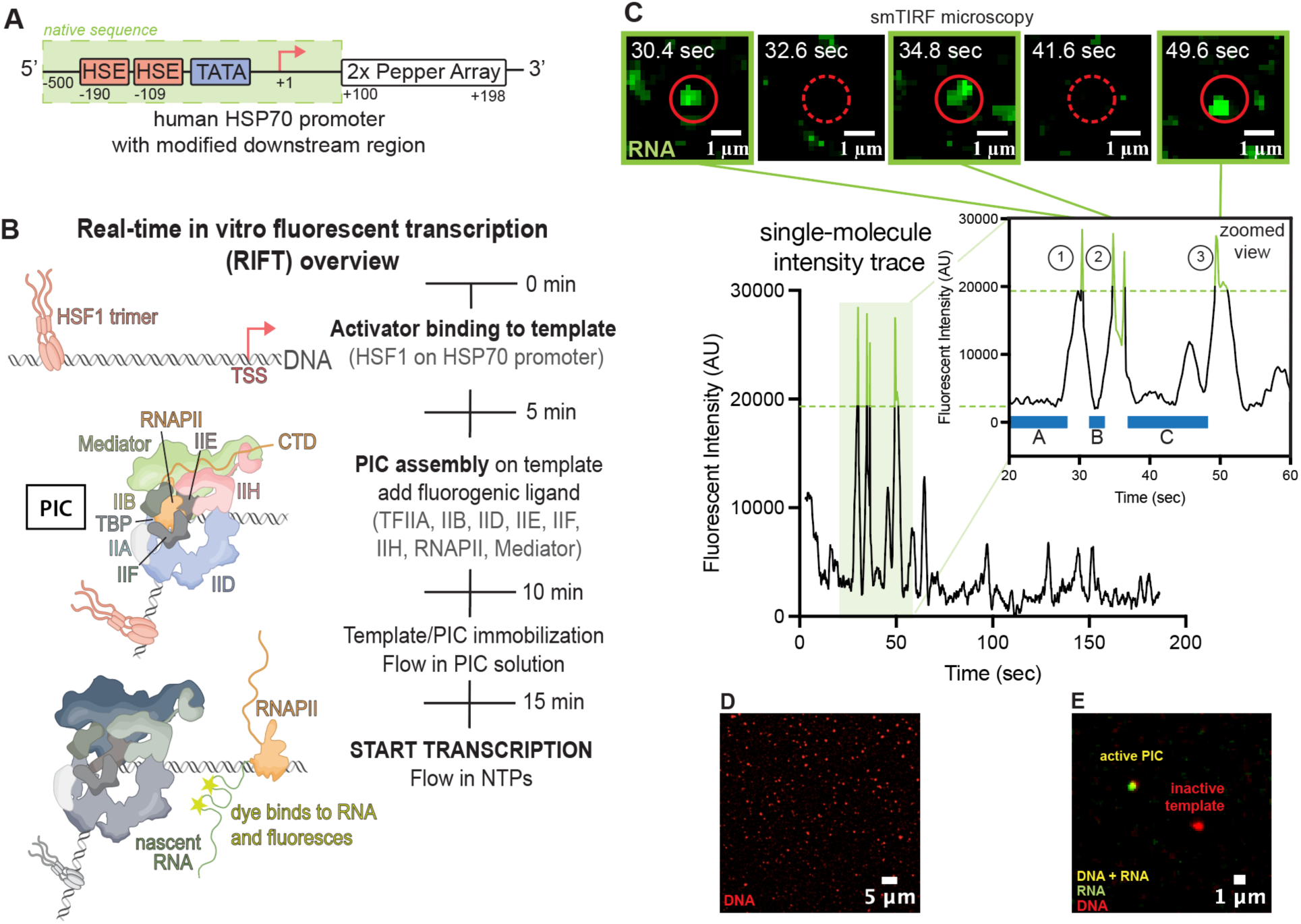
Real-time *in vitro* fluorescent transcription (A) Schematic of native HSPA1B promoter template with native sequence highlighted in green. (B) Overview of RIFT assay. (C) Representative RIFT images of RNA produced from immobilized HSPA1B promoter with corresponding intensity trace below. In this case, three transcripts were generated from this promoter. Blue bars illustrate (A) the time to first round transcription, (B) the time to second round transcription (i.e. re-initiation), and (C) is the time until another re-initiation event. Green dashed line represents signal filter which serves to distinguish RNA transcripts from background noise with high confidence. (D) Representative smTIRF image of immobilized HSPA1B DNA template labeled with Cy5 (see Methods). (E) Representative smTIRF image of immobilized promoters (red) during RIFT. In this case, an RNA (green) colocalized with DNA (yellow), indicative of an RNAPII transcript.

As shown in **Figure 1C** and **Video S1**, the RIFT assay allowed direct visualization of RNAPII transcription at individual promoters. Interestingly, it was immediately evident that re-initiation events were occurring at a subset of individual promoters (e.g. **Figure 1C**). Through direct labeling of the DNA templates (**Figure 1D**), we confirmed that ∼800 promoters were visualized in a single experiment simultaneously, and 100-200 templates typically generated at least one transcript within the 3min experimental timeframe (**Figure 1E**), yielding a template usage consistent with bulk *in vitro* studies,^47–49^ despite the shorter timeframes used here. Because all templates are identical, the number of active templates reflects the probability that RNAPII will successfully generate a “runoff” transcript.

Real-time measurement of RNAPII transcription at hundreds of active promoters allowed rigorous statistical analysis of population-level effects, to complement single-molecule resolution at individual promoters (**Figure S1C**). We confirmed that the signal from fluorophore-bound Peppers aptamers remained constant within the experimental timeframes, ruling out the possibility that intensity fluctuations simply resulted from photobleaching or transient compound dissociation from the RNA aptamer (**Figure S1D**). We also completed a series of control experiments i) to verify that spots represented single RNA transcripts, ii) to verify that transcript visualization was transient due to RNAPII “run off” from the template, and iii) to rigorously define signal vs. noise (**Figure S1D-F**). Furthermore, we completed negative control “drop out” experiments to verify that transcript detection required RNAPII (**Figure S1G**). Finally, we confirmed that RIFT demonstrated consistent and reproducible outcomes across multiple independent experiments (n=6; **Figure S1H**).

Based on the results summarized in **Figure 1** and **Figure S1**, we concluded that the RIFT assay could reliably measure human RNAPII transcriptional output at hundreds of promoters simultaneously, with high spatial and temporal resolution. We next shifted our focus to the mechanistic roles of individual PIC components, as well as the sequence-specific TF, HSF1.

### TFIIE & TFIIH increase transcription; TFIID accelerates first-round RNAPII activation

As a starting point for assessing PIC function, we tested RNAPII, TBP, TFIIB, and TFIIF, which is considered the “minimal system” because these four factors are sufficient for RNAPII transcription initiation *in vitro*.^5,6^ Separately, we tested PICs that included additional factors, to directly compare overall RNAPII transcriptional output. In each case, HSF1 was pre-bound to the HSPA1B promoter for PIC assembly. As expected, the minimal PIC exhibited reduced activity compared to PICs that additionally included TFIIE, TFIIH, and TFIIA (**Figure 2A**).^7,8^ Mediator further increased RNAPII transcription from TBP-containing PICs (**Figure 2A, B**). By contrast, TFIID-containing PICs attenuated overall transcriptional output vs. TBP-containing PICs in this assay (**Figure 2A, B; Figure S2A**). (Note that TFIID contains TBP plus 13 TBP-associated factors, called TAFs.) This result highlights the fact that RIFT detects transcripts during the “elongation” phase of RNAPII transcription (over 100bp downstream of the TSS; runoff transcription at +198), after the promoter-proximal pause region. Prior experiments have shown that TFIID enables RNAPII promoter-proximal pausing, whereas TBP-containing PICs do not pause.^46^ This provides a straightforward mechanistic explanation for reduced overall RNAPII transcripts with TFIID-containing PICs compared with TBP in the RIFT assays.

**Figure 2.**
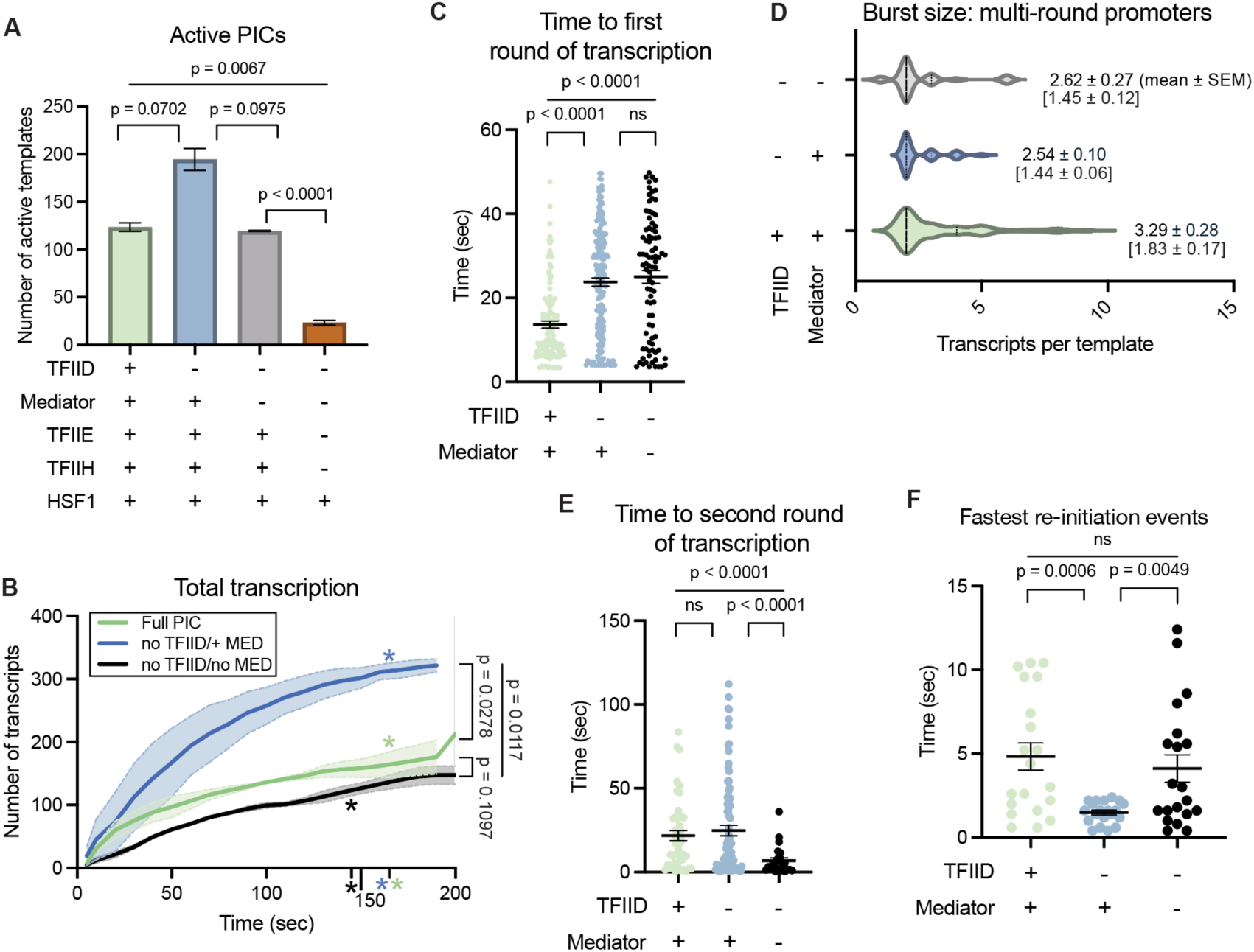
TFIIE and TFIIH increase transcription; TFIID accelerates RNAPII activation (A) Number of active promoters based upon PIC composition. All lanes include minimal system of TBP, TFIIB, TFIIF, and RNAPII; other PIC factors added as noted. If TFIID was added, TBP was not included in addition. Bars represent mean ± SEM. (B) Total transcription over time from PICs ±Mediator and/or TFIID. Asterisks signify burst duration, line depicts mean values, and shading represents SEM. No TFIID experiments included TBP instead. (C) Time to first round transcription from Full PIC (left) or PICs lacking TFIID or Mediator and TFIID. If no TFIID, TBP was used. Black bars represent mean ± SEM. (D) Burst size as function of Mediator and TFIID. Values listed are mean burst sizes from multi-round promoters (±SEM); values in brackets are mean burst size of all active promoters (±SEM). (E) Time to re-initiation with Full PIC (left) or PICs lacking TFIID or Mediator and TFIID. If no TFIID, TBP was used. Black bars represent mean ± SEM. (F) Fastest re-initiation events (n=20). Black bars represent mean ± SEM. All panels shown (A-F) represent data from 2 biological replicates.

Notably, PICs containing Mediator and TFIID (i.e. complete PICs) resulted in the fastest “first round” RNAPII activation (**Figure 2C; Figure S2B**), and also showed evidence for longer burst durations (green asterisks, **Figure 2B**) compared with PICs lacking Mediator and TFIID. Here, we define burst duration as the time post-NTP addition until the transcription rate becomes less than 1 every 6sec (see Methods). PICs lacking TFIID also showed long burst durations (**Figure 2B**), suggesting a role for Mediator (see Discussion). Note that our reported rate of first round RNAPII transcription considers only the first 60sec, to balance PIC-specific differences in burst duration. For example, few additional templates are activated after 60sec in the absence of Mediator and TFIID, in contrast with Mediator-containing PICs (**Figure S2B**).

We next determined whether TFIID and/or Mediator increased the the probability that a once-activated PIC would re-initiate. As shown in **Figure S2C**, the fraction of active templates that re-initiate was similar across conditions, whereas the average number of transcripts generated from multi-round promoters (i.e. burst size) increased with the full PIC (**Figure 2D**). Re-initiation rates were similar or slightly reduced with the “complete” PIC compared with PICs lacking TFIID and Mediator (**Figure 2E**). However, as with first-round transcription, we noticed that re-initiation occurred across longer timeframes with PICs containing TFIID and/or Mediator (**Figure 2E**), which skews the overall re-initiation rate compared with PICs lacking TFIID and Mediator. We therefore compared the fastest re-initiation events (n=20) for each condition (**Figure 2F**). Taken together, the data in **Figure 2E, F** showed that PICs containing Mediator and TFIID extend the timeframe for RNAPII re-initiation (i.e. increased burst duration) and that Mediator accelerated re-initiation rates.

### HSF1 and Mediator cooperatively activate RNAPII through multiple mechanisms

The data summarized in **Figure 2** and **Figure S2** confirmed basic roles for TFIIE, TFIIH, and TFIID and established benchmarks for RNAPII transcription activation with the RIFT system. We next focused on Mediator and the TF HSF1, based in part upon their paramount importance for RNAPII regulation in cells.^10,50^ Because Mediator and TFIID have been shown to coordinately regulate RNAPII transcription *in vitro* and in cells,^16,17,51^ all subsequent experiments included TFIID rather than TBP.

Compared with the full PIC, removal of both HSF1 and Mediator had dramatic negative effects on RNAPII transcription, greater in magnitude than loss of either factor alone. Fewer templates generated transcripts (**Figure 3A; Figure S3A**)—suggesting a lower probability of RNAPII activation—and no re-initiation occurred in the absence of HSF1 and Mediator (**Figure 3B**). Collectively, these defects yielded a massive reduction in overall RNAPII transcription (**Figure 3C**). Removal of HSF1 or Mediator individually had intermediate impacts on RNAPII transcription compared with loss of both factors at once (**Figure 3A-D; Figure S3B**). A notable exception was that only Mediator altered the proportion of active templates undergoing re-initiation (**Figure S3C**), suggesting a key role independent of HSF1 binding. Separate sets of RIFT experiments compared full PICs vs. PICs lacking either Mediator or HSF1, and the results are summarized in **Figure S4**.

**Figure 3.**
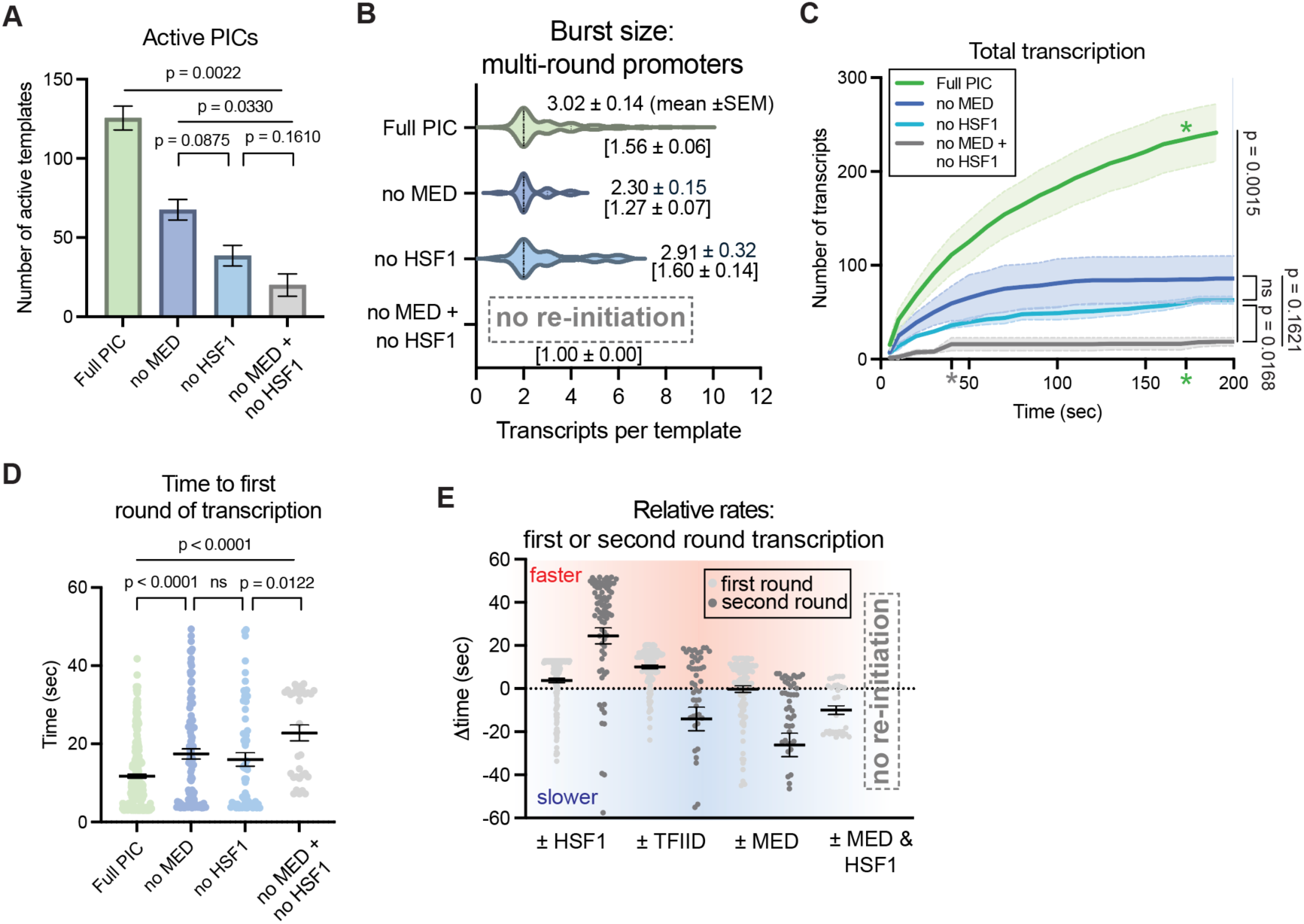
HSF1 and Mediator cooperatively activate RNAPII transcription (A) Number of active PICs ±HSF1, ±Mediator, or ±Mediator and HSF1. Bars represent mean ± SEM. (B) Burst size as function of Mediator and HSF1. Values listed are mean burst sizes from multi-round promoters (±SEM); values in brackets are mean burst size of all active promoters (±SEM). (C) Total transcription over time as a function of Mediator and HSF1. Asterisks signify burst duration, line depicts mean values, and shading represents SEM. (D) Time to first round transcription from Full PIC (left) or PICs lacking factors as indicated. Black bars represent mean ± SEM. (E) Summary of relative rates for 1st or 2nd round transcription from PICs ±HSF1, ±TFIID, ±Mediator, and ±Mediator and HSF1. The time difference reflects the impact of that factor compared with assays in its absence. Positive values (red shading) indicate faster rate; negative values indicate slower rate (blue shading). Black bars represent mean ± SEM. All panels shown (A-E) represent data from 2 biological replicates, except n=6 for Full PIC experiments.

Combined loss of Mediator and HSF1 also significantly reduced the rates of RNAPII activation within promoter-bound PICs. As shown in **Figure 3D**, generation of a “first round” transcript was significantly faster in the presence of Mediator and HSF1 (i.e. full PIC). Whereas re-initiation did not occur in the absence of Mediator and HSF1, comparison of full PIC re-initiation rates vs. removal of Mediator or HSF1 alone suggested that HSF1 has a dominant effect on RNAPII re-initiation rates (**Figure S3B**). Moreover, combined loss of Mediator and HSF1 markedly reduced the burst size (**Figure 3B**) and burst duration (gray asterisks, **Figure 3C**). For example, RNAPII activation effectively stopped within 1min NTP addition in the absence of Mediator and HSF1, whereas full PICs continued to activate throughout the duration of the experiment (3min; **Figure S3D**). A summary of how TFIID, HSF1 and/or Mediator impact RNAPII first-round transcription or re-initiation rates is shown in **Figure 3E**.

### Mediator accelerates HSF1 target search for rapid RNAPII activation

Like most sequence-specific, DNA-binding TFs, HSF1 has a modular architecture, with a DNA-binding domain (DBD; residues 1-221) and a disordered activation domain (AD; residues 407-529). Activation of heat shock response genes by HSF1 has shown a Mediator dependence in *Drosophila*,^52^ and HSF1 appears to support rapid RNAPII re-initiation and bursting under heat shock conditions.^22^ The native human HSPA1B promoter used in our RIFT assays has two HSF1 binding sites upstream of the TSS (**Figure 1A**).

HSF1 is a stimulus-response TF and under normal conditions HSF1 is inactive and sequestered in the nucleus and cytoplasm by the HSP70 chaperone. Upon heat shock stimulation, HSP70 dissociates and HSF1 accumulates in the nucleus, promoting DNA binding and target gene activation.^53^ To replicate this stimulus response, we modified the RIFT assay such that PICs were pre-assembled on promoters in the absence of HSF1. HSF1 was then introduced concurrently with NTPs. As expected, HSF1 activated RNAPII transcription under these conditions, similar to pre-bound HSF1 experiments (**Figure 4A; Figure S5A**). In fact, RNAPII activation occurred even faster under this “stimulus response” condition compared with standard RIFT assays with pre-bound HSF1 (**Figure 4B, C**).

**Figure 4.**
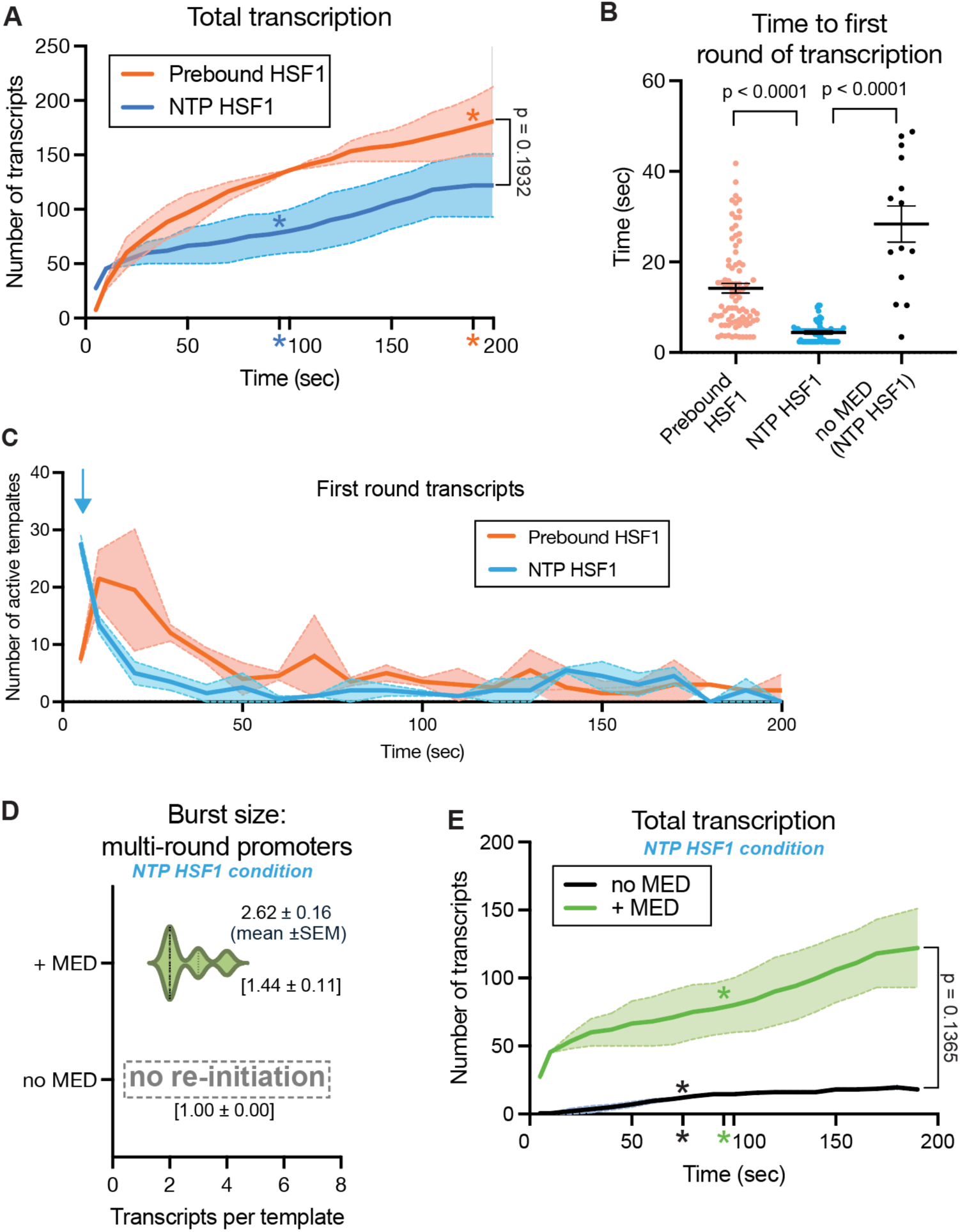
Mediator accelerates HSF1 target search for rapid RNAPII activation (A) Total transcription over time, comparing pre-bound HSF1 (standard condition) vs. HSF1 added with NTPs (stimulus response condition). Asterisks signify burst duration, line depicts mean values, and shading represents SEM. (B) Time to first round transcription comparing pre-bound HSF1 vs. HSF1 added with NTPs; also shown is data from NTP HSF1 condition in absence of Mediator. Black bars represent mean ± SEM. (C) Plot showing first round transcripts over time for pre-bound HSF1 vs. HSF1 added with NTPs. Blue arrow highlights faster RNAPII activation in NTP HSF1 condition. Line depicts mean values and shading represents SEM. (D) Burst size as function of Mediator in NTP HSF1 condition. Values listed are mean burst sizes from multi-round promoters (±SEM); values in brackets are mean burst size of all active promoters (±SEM). (E) Total transcription over time as a function of Mediator in NTP HSF1 condition. Asterisks signify burst duration, line depicts mean values, and shading represents SEM. All panels shown (A-E) represent data from 2 biological replicates.

Because HSF1 appears to activate RNAPII transcription through Mediator,^52^ we next tested whether the rapid RNAPII activation was Mediator-dependent. As shown in **Figure 4B**, Mediator substantially increased the rate of HSF1-dependent RNAPII activation compared with PICs assembled without Mediator. Furthermore, re-initiation was completely blocked in the absence of Mediator under the “stimulus response” condition (**Figure 4D; Figure S5B-D**). Consequently, overall RNAPII transcriptional output was substantially reduced in the absence of Mediator (**Figure 4E**), which was consistent with the pre-bound HSF1 experiments (**Figure S4I**).

Compared with “standard” RIFT assays with pre-bound HSF1, the Mediator-dependence was amplified under the stimulus response condition, such that re-initiation and rapid RNAPII activation did not occur in its absence. We consider the stimulus response experimental paradigm, in which HSF1 is added with NTPs, to better reflect physiological conditions compared with pre-bound HSF1 because NTPs are always available *in vivo*, including the moment HSF1 enters the nucleus and binds target gene promoters. Consequently, all remaining experiments were completed in this way.

### Mediator enables rapid TF-dependent RNAPII activation without TF-DNA binding

Mediator contains an unusually high percentage of intrinsically disordered regions (IDRs) among its 26 subunits.^39^ The HSF1 activation domain (AD) is also an IDR. A characteristic of IDRs is their large hydrodynamic radius; because they are unstructured, IDRs will rapidly sample a large number of conformational states. One way that IDRs may regulate transcription is by accelerating the “on rate” for factor binding because IDRs represent a much larger “target” compared with a compact, structured domain.^37^ We emphasize that this “IDR cloud” model isn’t mutually exclusive with molecular condensates, but it does not require condensate formation.

As shown in **Figure 4B, C**, HSF1 activation of RNAPII was so fast with the “stimulus response” experimental scheme, we wondered whether the HSF1 DNA-binding domain (DBD) was even required. Specifically, we hypothesized that rapid, Mediator-dependent activation could be occurring, at least in part, through IDR-IDR interactions between HSF1 and Mediator. We selected the AD from the TF SREBP to test this hypothesis, because it is among the best characterized TF-Mediator interactors.^54^ The SREBP-AD (residues 1-50) is an IDR and it is challenging to isolate on its own. Therefore, we tethered the SREBP-AD to glutathione-S-transferase (GST). We then completed “stimulus response” RIFT assays in which GST-SREBP-AD was added with NTPs.

Remarkably, GST-SREBP-AD activated RNAPII at least as well as HSF1 in these experiments (**Figure S6A**), with increased numbers of active templates (**Figure 5A**) and overall transcriptional output (**Figure 5B**). GST-SREBP-AD also increased RNAPII burst duration (**Figure 5B**) with modest effects on burst size (**Figure 5C**). GST-SREBP-AD also accelerated the rate of re-initiation (**Figure 5D; Figure S6B**), similar to HSF1. Interestingly, however, first-round RNAPII activation occurred more rapidly with HSF1 compared with GST-SREBP-AD (**Figure 5E**). In fact, the rate of first-round activation was unchanged ±GST-SREBP-AD, in contrast with ±HSF1 experiments (**Figure S4B**).

**Figure 5.**
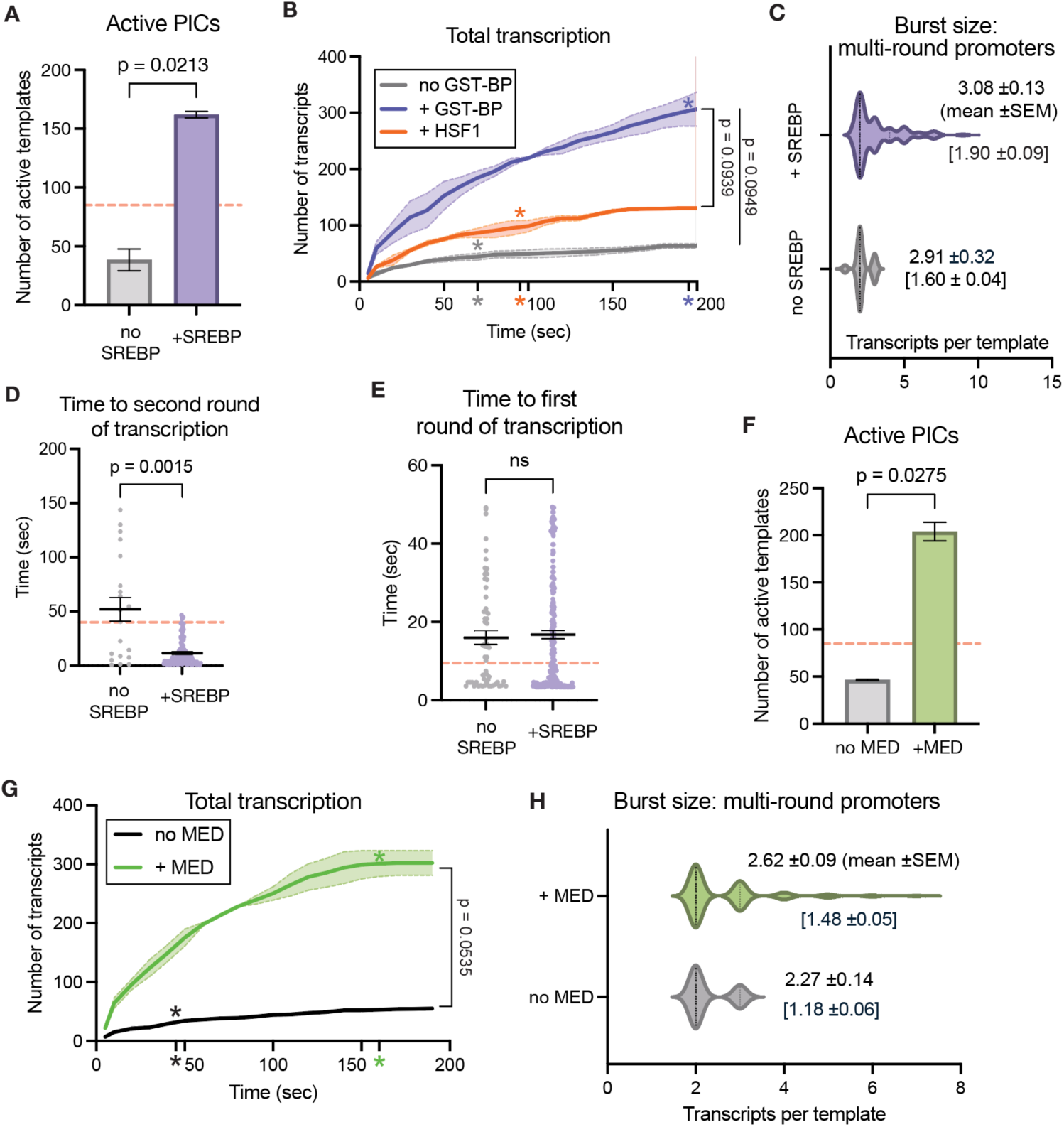
Mediator enables rapid TF-dependent RNAPII activation without TF-DNA binding (A) Number of active PICs ±GST-SREBP, in which GST-SREBP added with NTPs (i.e. stimulus response condition). For comparison, dashed orange line shows mean number of active PICs for NTP HSF1 condition. Bars represent mean ± SEM. (B) Total transcription over time, comparing ±GST-SREBP with NTP HSF1 or no GST-SREBP, all under stimulus response condition. Asterisks signify burst duration, line depicts mean values, and shading represents SEM. (C) Burst size as function of GST-SREBP. Values listed are mean burst sizes from multi-round promoters (±SEM); values in brackets are mean burst size of all active promoters (±SEM). (D) Time to re-initiation ±GST-SREBP. Dashed orange line conveys NTP HSF1 mean time to re-initiation. Black bars represent mean ± SEM. (E) Time to first round transcription as a function GST-SREBP. For comparison, dashed orange line depicts data from NTP HSF1 experiments. Black bars represent mean ± SEM. (F) Number of active PICs ±Mediator, in which GST-SREBP added with NTPs (i.e. stimulus response condition). For comparison, dashed orange line shows mean number of active PICs for NTP HSF1 condition. Bars represent mean ± SEM. (G) Total transcription over time ±Mediator in NTP GST-SREBP condition. Asterisks signify burst duration, line depicts mean values, and shading represents SEM. (H) Burst size as function of Mediator in NTP GST-SREBP condition. Values listed are mean burst sizes from multi-round promoters (±SEM); values in brackets are mean burst size of all active promoters (±SEM). All panels shown (A-H) represent data from 2 biological replicates.

The SREBP-AD binds Mediator with high affinity,^54^ and prior reconstituted transcription assays showed that SREBP-Mediator cooperatively activate RNAPII transcription.^55,56^ Consequently, we expected that the activity of GST-SREBP-AD would require Mediator, and this was confirmed in RIFT experiments ±Mediator. As shown in **Figure 5F-H**, fewer PICs were activated, overall transcriptional output was reduced, and fewer re-initiation events occurred from active PICs in the absence of Mediator (**Figure S6C**). Experiments with fluorescently labeled GST-SREBP-AD confirmed that Mediator increased GST-SREBP-AD dwell time on the HSPA1B promoter (**Figure S6D**), consistent with biochemical and structural data that showed direct SREBP-Mediator interaction.^54^ Notably, RNAPII “first round” activation rates were unchanged with GST-SREBP-AD ±Mediator under the stimulus response experimental conditions (**Figure S6E**), in contrast to similar experiments with HSF1 (**Figure 4B**). Moreover, Mediator accelerated re-initiation rates with GST-SREBP-AD, but only when comparing the fastest events (**Figure S5F**). Mediator increased the timeframe for re-initiation to occur, therefore, Mediator-dependent effects on re-initiation rates were negated across the entire population (**Figure S5G**).

Because the SREBP-AD is an IDR, we next tested whether its activity in the RIFT assays might involve condensate formation. We fluorescently labeled GST-SREBP-AD to directly visualize; however, at concentrations used in RIFT experiments (13nM), no evidence for condensates or clusters was observed (**Figure S5H, Video S2**). The spot sizes were consistent only with single molecules.

Collectively, the results summarized in **Figure 5** and **Figure S6** demonstrated that i) a DBD was not required for rapid, TF-dependent RNAPII activation and that ii) Mediator was sufficient for TF recruitment to the PIC, via TF AD-Mediator interactions. Furthermore, the RIFT experiments showed that iii) condensate formation did not contribute to SREBP-AD function, but SREBP-dependent PIC activation required Mediator.

### MED1-IDR can replace HSF1 to activate RNAPII transcription

To build from the GST-SREBP-AD results, and to further probe the condensate or “IDR cloud” models, we tested whether an IDR alone could functionally replace the HSF1 AD. To address this question, we cloned, expressed, and purified MED1-IDR tethered to the native HSF1 DBD (DBD-MED1-IDR; **Figure 6A**). Importantly, the HSF1 AD was removed and replaced with the MED1-IDR. The MED1-IDR is a well-tested domain that forms condensates *in vitro*.^57,58^

**Figure 6.**
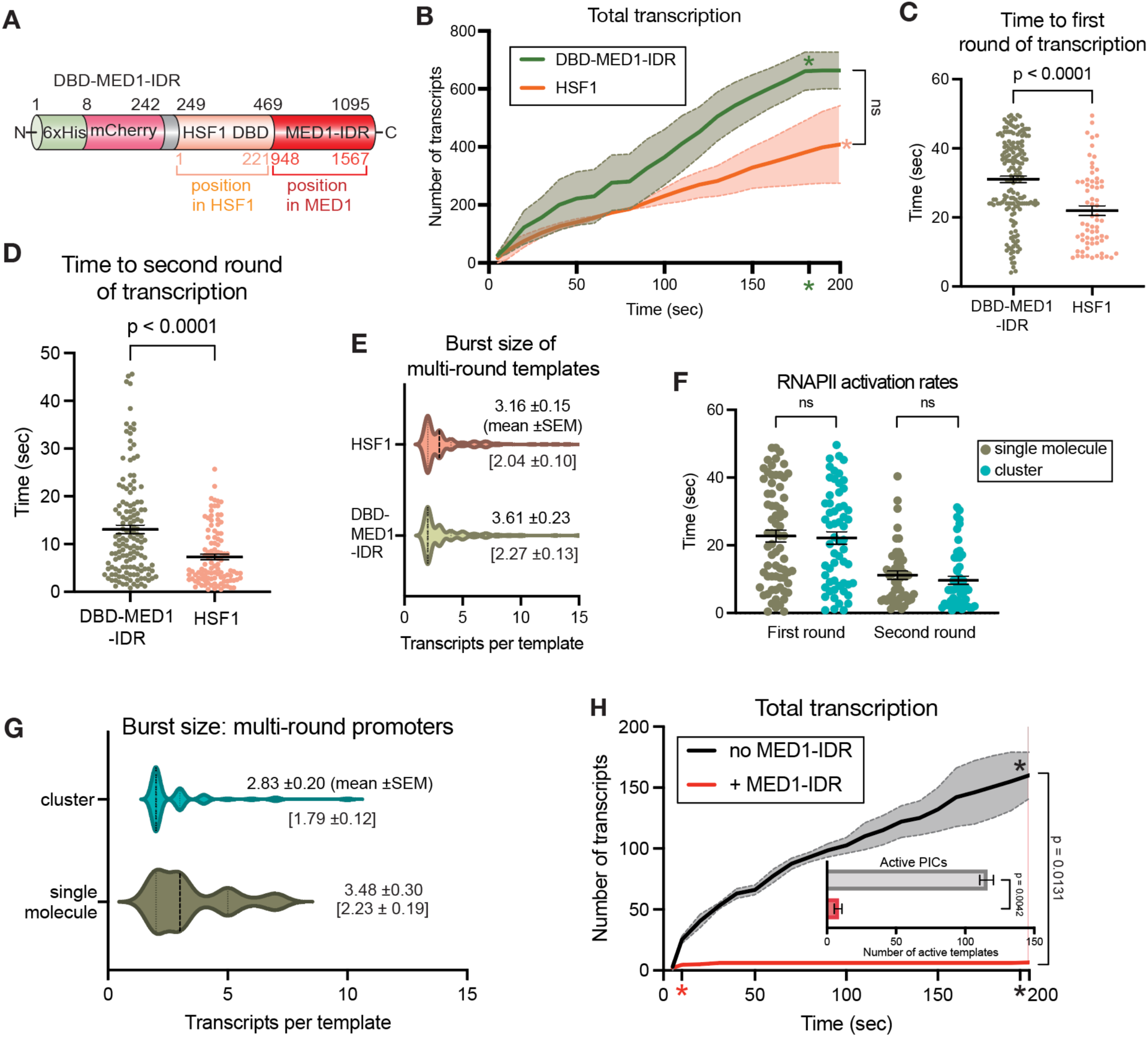
MED1-IDR can replace HSF1 to activate RNAPII activation and re-initiation (A) Schematic of DBD-MED1-IDR. (B) Total transcription over time for DBD-MED1-IDR or HSF1 (stimulus response conditions). Asterisks signify burst duration, line depicts mean values, and shading represents SEM. (C) Time to first round transcription for DBD-MED1-IDR or HSF1 experiments. Black bars represent mean ± SEM. (D) Time to re-initiation for DBD-MED1-IDR or HSF1 experiments. Black bars represent mean ± SEM. (E) Burst size from PICs with DBD-MED1-IDR or HSF1. Values listed are mean burst sizes from multi-round promoters (±SEM); values in brackets are mean burst size of all active promoters (±SEM). (F) Time to first- or second-round RNAPII transcription comparing single DBD-MED1-IDR molecules versus clusters. Black bars represent mean ± SEM. (G) Burst sizes from promoters associated with single DBD-MED1-IDR molecules versus clusters. Values listed are mean burst sizes from multi-round promoters (±SEM); values in brackets are mean burst size of all active promoters (±SEM). (H) Total transcription over time ±MED1-IDR (free; not DBD tethered). Asterisks signify burst duration, line depicts mean values, and shading represents SEM. Inset: number of active PICs ±MED1-IDR. Bars represent mean ± SEM. Panels B-H represent data from 2 biological replicates.

To enforce localization of the MED1-IDR to the HSPA1B promoter, we generated a modified HSPA1B template in which 17 consensus HSF1 binding sites were inserted upstream of the native HSF1 site (**Figure S7A**). Thus, this modified HSPA1B promoter contained 19 HSF1 sites compared with 2 for the native promoter. Note that the core promoter sequence remained unchanged (i.e. unchanged from the HSE at −109 to the beginning of the 2X Peppers array at +100).

We next used the “19X” template to test whether forced localization of the MED1-IDR might “super-activate” RNAPII transcription from the HSPA1B promoter. The DBD-MED1-IDR chimera was tested in a series of RIFT assays, in comparison with WT HSF1, and their concentrations were increased to 100nM in these experiments (HSF1 13nM previously) to better assess TF-dependent effects. Remarkably, the DBD-MED1-IDR was able to activate RNAPII similarly to WT HSF1, despite lacking the native HSF1 AD (**Figure 6B; Figure S7B**). Interestingly, however, WT HSF1 acted with faster kinetics for PIC first round activation (**Figure 6C**), and was capable of faster second round re-initiation (**Figure 6D; Figure S7C**), although each protein generated similar burst sizes from multi-round templates (**Figure 6E**). HSF1 had longer dwell times compared with DBD-MED1-IDR (**Figure S7D**), which may contribute to the faster RNAPII activation rates.

Confocal microscopy with the DBD-MED1-IDR chimera (100nM) confirmed that large clusters/condensates formed in the absence of immobilized DNA templates (**Figure S7E**). In RIFT assays, we also observed DBD-MED1-IDR clusters, in addition to single proteins. However, nearly all (96%; **Figure S7F**) DBD-MED1-IDR clusters comprised only 2-3 proteins and thus do not represent condensates. We next sorted DBD-MED1-IDR clusters from single molecules and linked their occupancy to subsequent RNAPII transcription from individual promoters. In this way, we could evaluate whether DBD-MED1-IDR clusters were functionally distinct from single molecules in the same experiment. As shown in **Figure 6F, G**, the DBD-MED1-IDR clusters did not increase RNAPII transcriptional output beyond that achieved by single molecules.

Taken together, these results demonstrated that i) the MED1-IDR can functionally replace a TF AD, invoking new mechanisms for RNAPII activation within the PIC; ii) WT HSF1 could more rapidly activate RNAPII compared with DBD-MED1-IDR, and iii) MED1-IDR clusters did not “super-activate” RNAPII compared with single molecules.

### Free MED1-IDR forms condensates, sequesters RNAPII, but squelches transcription

We next wondered about the mechanism by which the DBD-MED1-IDR activated RNAPII transcription from the 19X HSPA1B promoter. Based upon prior experiments showing that MED1-IDR partitions with RNAPII,^58,59^ we hypothesized that DBD-MED1-IDR might activate PICs by recruiting RNAPII. To test this concept, we expressed and purified an mCherry-tagged MED1-IDR (MED1 residues 948-1567). Using confocal microscopy, we confirmed that the MED1-IDR formed clusters/condensates (**Figure S7G**) and sequestered RNAPII (**Figure S7H**), in agreement with prior reports.^58,59^ A larger average size for MED1-IDR clusters in the presence of RNAPII is consistent with condensate behavior.^34^

Given that free MED1-IDR formed condensates and partitioned with RNAPII, we hypothesized that MED1-IDR condensates might localize to promoter-bound PICs even in the absence of a tethered DBD, to activate RNAPII transcription. However, this was not observed. Few MED1-IDR condensates or clusters were captured at or near promoter DNA (**Video S3**).

Furthermore, RIFT assays with added MED1-IDR completely inhibited RNAPII transcription (**Figure 6H**). These results are consistent with a squelching effect seen in prior biochemical assays with nuclear extracts.^58^ The stark functional differences between DBD-MED1-IDR and freely diffusing MED1-IDR demonstrates that the MED1-IDR can activate RNAPII transcription as a single protein or cluster, but squelches transcription as a condensate.

### Increased HSF1 promoter occupancy drives RNAPII burst size and burst duration

Finally, we used RIFT to test the mechanistic links between TF dwell times and RNAPII burst size, which have been widely reported in yeast and mammalian cells.^60–62^ The underlying mechanisms remain unclear, in part because TFs interact with many transcriptional regulatory proteins, including Mediator but also chromatin remodeling and chromatin modifying complexes.^63,64^

We measured HSF1 dwell times and as expected, dwell times increased at the 19X promoter compared with the native HSPA1B promoter (**Figure S7I**), providing a “model system” to test whether the PIC factors were sufficient to recapitulate TF-dependent changes in RNAPII bursting, as seen in cells. As shown in **Figure 7A, B**, the 19X HSPA1B promoter markedly increased RNAPII transcriptional output. Mechanistically, this was due to i) increased probability of RNAPII activation, resulting in more active PICs (**Figure 7A; Figure S7J**), ii) increased burst size across all active promoters (**Figure 7C**) and iii) increased probability that active PICs would re-initiate (**Figure S7K**). Moreover, the 19X promoter increased RNAPII re-initiation rates (**Figure S7L**); however, first round RNAPII transcription was actually slower from the 19X promoter (inset, **Figure 7J**).

**Figure 7.**
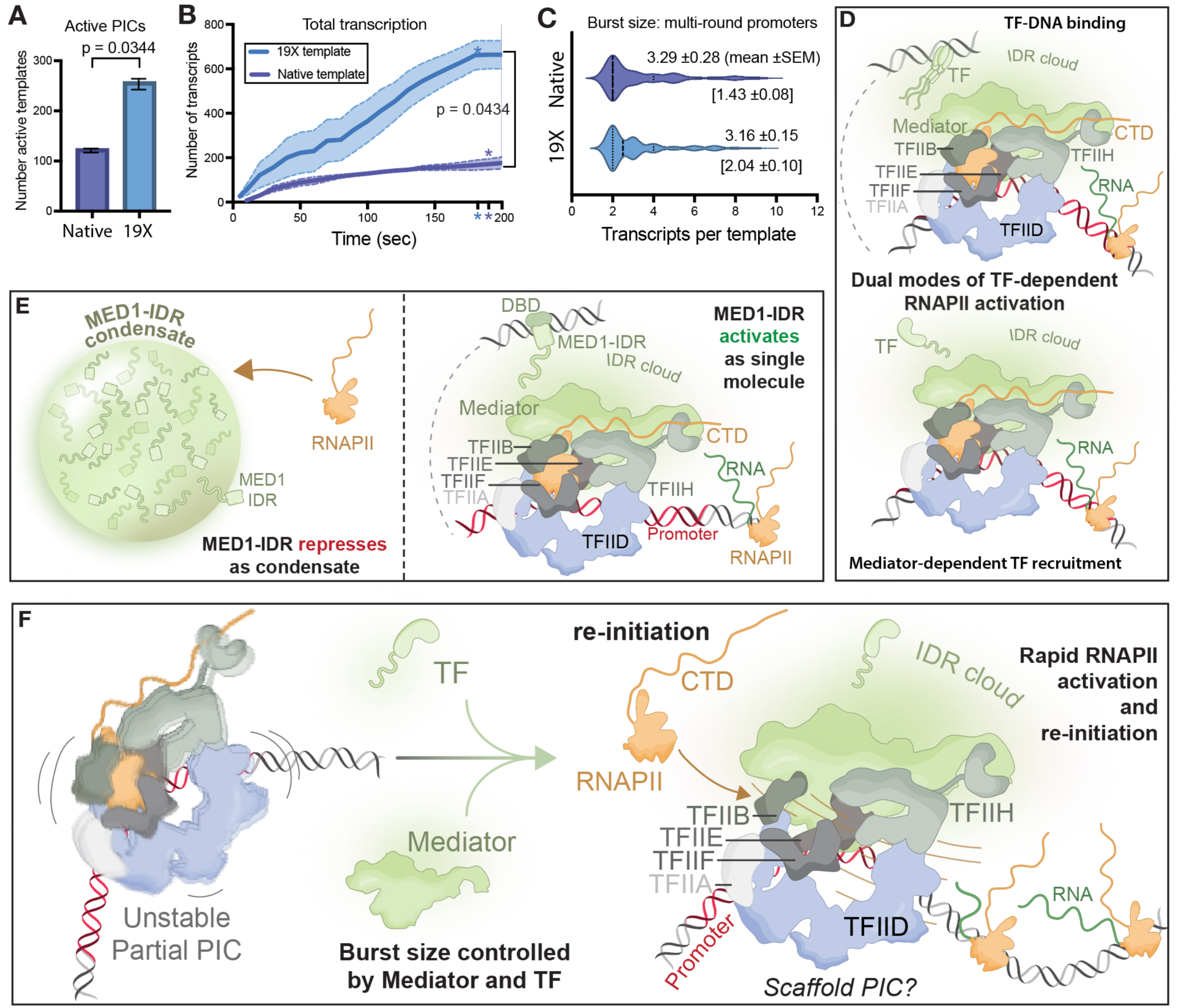
Increased HSF1 promoter occupancy drives RNAPII burst size and burst duration (A) Number of active PICs with native HSPA1B or 19X HSPA1B template. Bars represent mean ± SEM. (A) Total transcription over time, comparing native vs. 19X promoter templates. Asterisks signify burst duration, line depicts mean values, and shading represents SEM. (A) Burst size of native or 19X template. Values listed are mean burst size of multi-round promoters (±SEM) and values in brackets are mean burst size of all active promoters (±SEM). Panels A-C represent data from 2 biological replicates. (A) Model depicting two different TF-dependent activation mechanisms, one involving TF binding to promoter DNA (top), and the other that does not require DNA binding but instead the TF is recruited via Mediator (bottom). In each case, TF-Mediator interaction drives RNAPII activation. (A) The MED1-IDR adopts contrasting biochemical functions based upon its biophysical state. As a condensate, MED1-IDR squelches transcription, likely through sequestering RNAPII (left). As a single molecule, the MED1-IDR activates RNAPII transcription, likely through recruitment of RNAPII to the promoter. (A) Model depicting how TFs and Mediator enable rapid RNAPII activation and increase burst size and duration, likely through PIC stabilization and potentially stabilization of a PIC scaffold re-initiation complex. Mediator also increases the probability of RNAPII re-initiation after first-round transcription.

Collectively, these data are consistent with live cell imaging results that show TF-dependent control of RNAPII burst size and burst duration^60–62^ and underscore the paramount importance of TFs in controlling RNAPII function within the PIC. Our results with the 19X template further suggest that cellular strategies to increase TF-promoter occupancy are essential for high-level activation.

Because our RIFT experiments include only HSF1 and the PIC factors, our results establish that TF-PIC interactions, presumably through Mediator, contribute to TF-dependent regulation of RNAPII bursting either partially or entirely.

## Discussion

The insights from RIFT complement results obtained from live cell imaging and single-cell RNA-seq experiments, which only indirectly address mechanistic questions. Because RIFT combines smTIRF with reconstitution of human RNAPII transcription, we could directly assess molecular mechanisms and identify the contributions of specific factors. RIFT allows second-by-second visualization of RNAPII transcription across a population of hundreds of promoters at once, and direct visualization of RNAPII re-initiation from individual promoters. Furthermore, the RIFT system provided a means to distinguish single TF proteins from clusters or condensates, to determine whether condensates *per se* were required for rapid TF-dependent PIC activation, RNAPII re-initiation, or other processes. Each of these regulatory events, and their rates, are fundamental for RNAPII-dependent gene expression, but have not been directly visualized in a defined, biochemically reconstituted system before, to our knowledge.

The results from the RIFT assays yielded mechanistic insights that help reconcile confounding and/or unsettled models derived from live cell imaging experiments. We summarize in **Figure 7D-F** and provide some additional description in the following paragraphs.

### Enhancer-promoter communication

One of the most important but confounding questions in the transcription field is how enhancers regulate RNAPII activity at promoters.^65^ A straightforward model involves direct E-P contact, in which enhancer-bound TFs activate a promoter-bound PIC (**Figure 7**). Live cell imaging experiments suggested an alternative model that is not mutually exclusive, involving enhancer “activation at a distance”, via a factor(s) that dissociates from the enhancer to a nearby promoter.^10,66^ An abundance of evidence supports the “activation at a distance” model;^24–26^ however, it remains speculative because of the inherent limitations of live cell imaging. For example, the spatial resolution is not sufficient to distinguish direct vs. indirect E-P interactions.^19,35^ Moreover, because E-P interactions are transient and asynchronous across cells, fluorescence microscopy may not capture such events due to limits in temporal resolution.^36,67^

Our RIFT experiments provided insights about potential mechanisms for enhancer-dependent activation at a distance. Analysis of HSF1 under “stimulus response” conditions indicated that rapid TF-dependent RNAPII activation occurs in a Mediator-dependent manner. More importantly, we demonstrate that a TF activation domain without a DBD (i.e. GST-SREBP-AD) is sufficient for rapid RNAPII activation, but requires a Mediator-bound PIC. In fact, we demonstrate that Mediator is sufficient for TF recruitment to the PIC. This result suggests a straightforward mechanism in which a TF could diffuse from an enhancer to activate a nearby promoter via TF-Mediator binding (**Figure 7D**), and would not even require concomitant DNA-binding to the promoter. In cells, it is not possible to prove that TF recruitment to specific genomic loci can be DBD-independent but Mediator-dependent; however, supporting evidence derives from live cell imaging, in which the TF AD and DBD were each shown to influence residence times or recruitment/clustering on genomic DNA.^68,69^ We hypothesize that the Mediator “IDR cloud” contributes to TF recruitment in cell nuclei, providing a means by which IDR-rich TFs could cluster without requiring phase separation. Finally, we emphasize that a TF-PIC recruitment mechanism that circumvents the DBD has significant biological consequences, given the emerging evidence that TF isoforms lacking the DBD are widely expressed in cell-type and context-specific ways.^70^

### Molecular condensates and IDR-dependent RNAPII regulation

In light of recent evidence suggesting that condensates regulate RNAPII function in cells, we were surprised to discover that rapid RNAPII activation kinetics occurred without condensate formation (**Figure 7E**). IDR-containing proteins can spontaneously phase separate, and HSF1, Mediator, and RNAPII have each been shown to phase separate under appropriate conditions *in vitro*.^57,59,71,72^ Phase separation via the formation of molecular condensates may have evolved to compartmentalize biological processes such as transcription initiation.^31,34^ A consequence is enforced high local concentrations of factors such as Mediator, TFs, and RNAPII.

For practical reasons, our RIFT assays included concentrations of Mediator, RNAPII, and HSF1 that were lower than required to form condensates *in vitro*, and their co-localization on promoter DNA did not induce condensate formation. However, low-affinity IDR-IDR interactions can be biologically meaningful in the absence of phase separation, even at low concentrations. For example, because they are unstructured, IDRs have a large hydrodynamic radius, which presents an “IDR cloud” that may accelerate on-rates with other IDR-containing proteins.^37^ In agreement, the DBD-MED1-IDR accelerated RNAPII re-initiation compared with experiments lacking HSF1 (compare **Figure 6D** & **Figure S5B**). Mediator contains an unusually large number of IDRs,^39^ and we observed that HSF1 RNAPII activation was significantly accelerated in the presence of Mediator, despite HSF1 having a DBD that can independently bind the promoter. A Mediator requirement for RNAPII activation was also observed with GST-SREBP-AD. However, because Mediator serves multiple roles in RNAPII activation, we cannot de-couple from IDR-specific effects.

Because RIFT can simultaneously visualize condensates and RNAPII transcription from single promoters, we were able to directly measure functional differences between proteins in their condensate or soluble forms. We show that rapid RNAPII activation occurs in the absence of condensate formation, but provide evidence that IDRs contribute to RNAPII transcriptional output in meaningful ways. For the MED1-IDR, we demonstrate that it squelches transcription as a condensate, but activates transcription as a soluble protein or cluster. Notably, MED1-IDR-dependent activation required tethering to promoter DNA, which may prevent condensate formation, based upon recent results with the Gcn4 TF in yeast.^73^

Our results are most consistent with reports that suggest minor or specialized roles for condensates in the regulation of gene expression in cells.^69,74,75^ Drawing parallels from the “sub-optimization” of TF binding affinities on genomic DNA,^76,77^ we speculate that similar sub-optimization evolved among IDR-rich transcription regulatory proteins, such that their normal biological function typically does not require condensate formation. Consistent with this idea, TFs designed with enhanced condensate-forming properties had “gain-of-function” activity and reduced target gene specificity.^78^ We plan to further address the role of condensates in future projects. However, recent work from Bremer et al. suggests that potential condensate-dependent RNAPII regulatory mechanisms will not be intuitive or straightforward.^73^

Contrary to expectations, we showed that the MED1-IDR was sufficient for PIC activation and can functionally replace a TF activation domain. Because the MED1-IDR can sequester TFs and RNAPII,^57–59,79^ we hypothesize that the IDR alone—if localized to the promoter—enhances PIC assembly and stability. However, the kinetics of first-round PIC activation were slower with the DBD-MED1-IDR compared with WT HSF1. We speculate that faster first-round PIC activation with WT HSF1 reflects a requirement for direct HSF1 AD-Mediator interaction, to trigger conformational shifts that activate RNAPII. Prior work has yielded evidence for large-scale structural re-organization of human or yeast Mediator upon TF binding,^80^ including with the SREBP-AD,^56^ and TF-dependent structural shifts correlate with RNAPII activation within the PIC.^81^ Potentially, Mediator structural isomerization occurs rapidly with WT HSF1, but more slowly with an artificial TF such as the MED1-IDR. Given the unstable nature of the PIC,^27,41^ this kinetic difference might have meaningful consequences in cells.

### RNAPII bursting and rapid activation coordinated by TF-Mediator within PIC

The PIC appears to be highly unstable in yeast and mammalian cells.^82^ In yeast, the PIC may persist on genomic DNA for 4-9sec,^27^ whereas RNAPII or Mediator clusters have a lifetime of approximately 10sec in mouse ESCs.^29,30^ These findings suggest that RNAPII initiation mechanisms evolved for rapid activation. Here, we demonstrate that PIC activation is extremely rapid under two different experimental conditions: a “standard” assay with HSF1 pre-bound for NTP addition, and a “stimulus response” assay in which HSF1 was added with NTPs. We identify the TF HSF1, TFIID, and Mediator as essential for fast first-round RNAPII activation.

Consequently, TFs, Mediator, and/or TFIID may serve as “gatekeepers” to prevent or minimize unregulated or random RNAPII initiation events that could otherwise disrupt normal transcriptional responses. In agreement, Mediator and TFs are generally linked to burst size regulation in cells,^23,62,83^ and the TAF1 subunit of TFIID was shown to limit RNAPII burst size during stimulus response in *Drosophila*.^84^

Interestingly, we observed that burst size and burst duration increased in a Mediator- and TF-dependent manner (**Figure 7F**), under both experimental paradigms (standard and stimulus response). These results suggest that TFs and Mediator stabilize the PIC and potentially a scaffold PIC (see below); Mediator also increased the probability that an active promoter would re-initiate, suggesting additional uncharacterized regulatory functions. Prior biochemical experiments have shown that TFs and Mediator stabilize PICs^85,86^ and have suggested that TF-Mediator interactions promote re-initiation.^86^ Because the PIC is inherently unstable *in vivo*,^27^ PIC stabilization could have meaningful biological consequences.

Data from live cell imaging experiments suggest that burst size and duration is Mediator-dependent^23,83^ and tracks with TF occupancy/residence time at promoters,^60–62^ in support of our RIFT data. To further probe this question, we generated an artificial HSPA1B template with 19 HSF1 binding sites. Consistent with cell-based results, this “19X” template i) increased HSF1 dwell time at the HSPA1B promoter, ii) increased burst size, iii) accelerated RNAPII re-initiation rates, and iv) increased the probability of re-initiation. Because RIFT uses a defined set of factors, our results demonstrate the PIC is sufficient to recapitulate these TF-dependent effects, and auxiliary factors are not required. In fact, burst sizes measured with RIFT on the native HSPA1B promoter are consistent with estimates from live-cell imaging and single-cell transcriptomics experiments.^87–91^ However, other factors present in cells likely contribute to burst size or duration through alternate or related mechanisms. For example, MYC, BRD4, AFF4, DSIF, or the PAF complex have also been shown to influence RNAPII bursting^19,90,92,93^ but the mechanistic basis remains unclear.

Although first-round RNAPII activation was fast, our RIFT data showed that re-initiation was even faster. In fact, we commonly observed re-initiation within 1 second in the presence of Mediator and HSF1. This rapid rate suggests that re-initiation occurs via a PIC scaffold complex,^86^ but its composition and potential mechanism of action remains unclear. One model consistent with the rapid kinetics is a single-stranded “DNA bubble” may briefly remain after RNAPII promoter escape, perhaps stabilized by a PIC factor. In this way, RNAPII could rapidly engage with ssDNA^6^ to re-initiate transcription, circumventing other rate-limiting steps required for *de novo* initiation.

Others have shown that *de novo* PIC assembly is rate-limiting for RNAPII activation *in vitro*.^17,94,95^ Our RIFT assays were designed such that PIC assembly occurred prior to NTP addition. The fact that re-initiation was still faster overall, compared with first-round transcription from pre-assembled PICs, further supports a scaffold model. That said, we acknowledge that the scaffold PIC remains controversial,^96^ poorly understood, and may function in a variety of compositional or structural states.^14,15,97^

### Limitations of this study

The reconstituted system we use for the RIFT assays does not match the complex array of proteins, nucleic acids, and metabolites that converge on an active gene in a human cell. This has advantages for addressing mechanistic questions but contributions from other regulatory factors cannot be assessed. In cells, transcription occurs on chromatin templates, whereas “nucleosome free regions” were used here. The concordance of our results with data derived from cell-based methods is striking and suggests that the PIC is the primary regulator of RNAPII function at gene 5’-ends in cells. The Peppers aptamer sequences were located beyond +100; consequently, we cannot measure promoter-proximal transcription and promoter-proximal paused transcripts will not be detected. We cannot exclude the possibility that under “Full PIC” conditions, some transcripts may be generated from partial PIC assemblies in the population. Finally, our HSPA1B promoter templates extended only to +198, which is after the promoter-proximal pause site, but does not allow a reliable assessment of RNAPII elongation.

## RESOURCE AVAILABILITY

Lead contact: Further information and requests should be directed to and will be fulfilled by the lead contact, Dylan Taatjes.

Material availability: Plasmids or reagents generated in this study are available from the lead contact upon request.

## Data and code availability

- All data reported in this paper will be shared by the lead contact upon request.
- All original code has been deposited at our Github repository and is publicly available at https://github.com/meganpalacio/RIFTA.
- Any additional information required to reanalyze the data reported in this paper is available from the lead contact upon request.

## Acknowledgments

We thank Bede Portz and Ann Boija (Dewpoint Therapeutics) for helpful advice; Jim Goodrich, Stephen Archuleta, and Joe Dragavon (UC-Boulder) for assistance with development of the RIFT system, and Theresa Nahreini (UC-Boulder) for cell culture assistance, Annette Erbse (UC-Boulder) for instrumentation support, and Kent Ritchie for development of the RIFTA script. We thank Jim Goodrich (UC-Boulder) for providing monoclonal TAF4 antibodies, and R. Tjian (UC-Berkeley) for monoclonal ERCC3 antibodies. RIFT imaging was performed at the UC-Boulder BioFrontiers Institute’s Advanced Light Microscopy Core (RRID: SCR_018302). TIRF microscopy was performed on a Nikon Ti2-E microscope supported by the Howard Hughes Medical Institute. Spinning disc confocal microscopy was performed on a Nikon Ti-E microscope supported by the BioFrontiers Institute and the Howard Hughes Medical Institute. We acknowledge the Shared Instruments Pool (RRID: SCR_018986) for the Department of Biochemistry at UC-Boulder for providing access to the Typhoon 5 Imager, funded by the NIH (S10OD034218). This work was supported by the NIH (R35 GM139550 to DJT; T32 GM065103 to MP) and the NSF (DGE-204034 to MP). Funding support was also provided by Dewpoint Therapeutics to DJT. We acknowledge funding support for instrumentation (S10OD025267).

## Author contributions

Experimental design: MP, DJT

Data collection and analysis: MP, DJT Supervision and funding: DJT

Writing: MP, DJT

## Supplemental information

Document S1. Figures S1-S7

**Video S1.** Real-time *in vitro* fluorescent transcription (related to Figure 1)

**Video S2.** Fluorescent GST-SREBP-AD during RIFT (related to Figure 5)

**Video S3.** mCherry-MED1-IDR during RIFT (related to Figure 6)

## STAR METHODS

### KEY RESOURCES TABLE

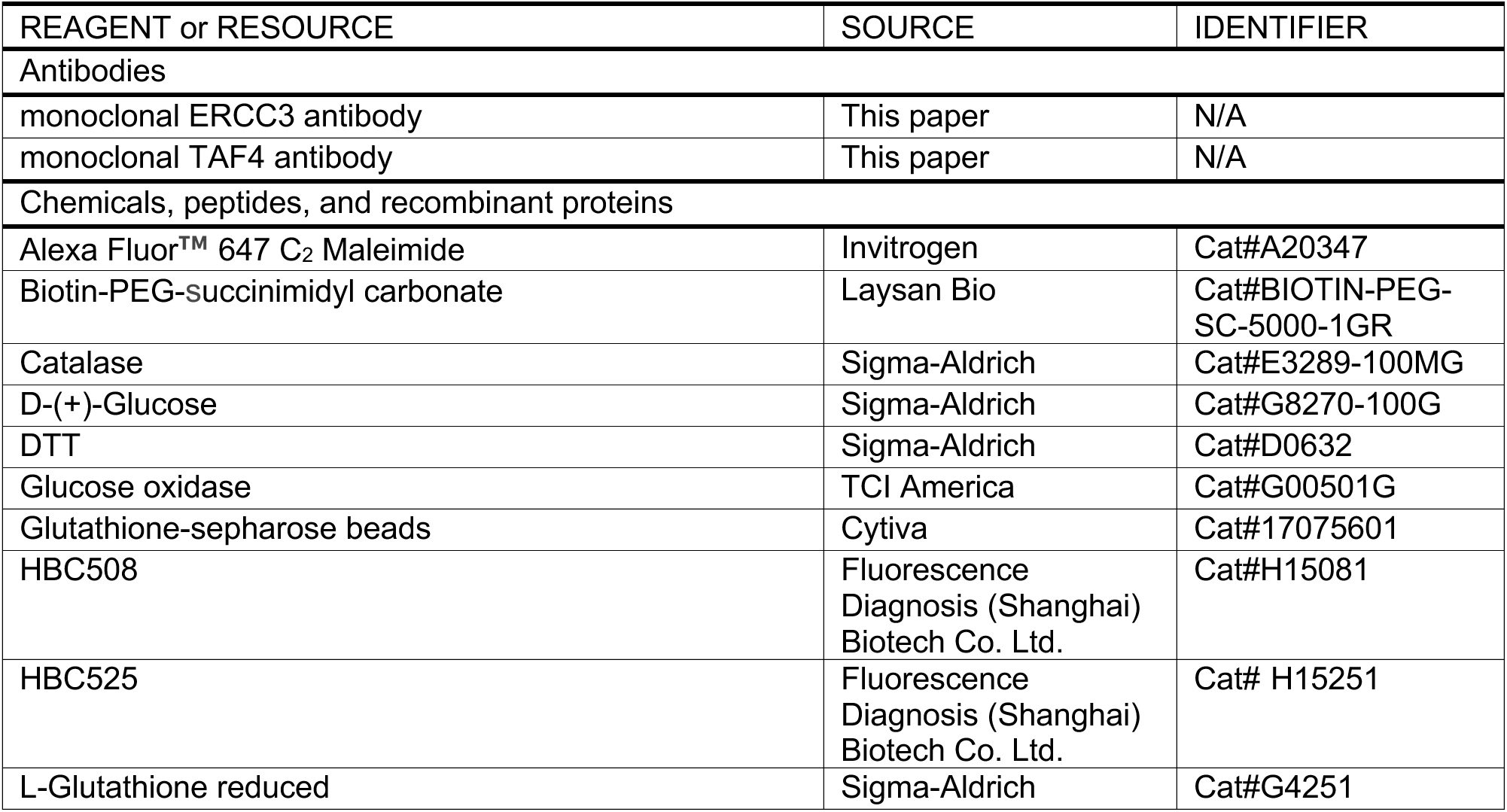
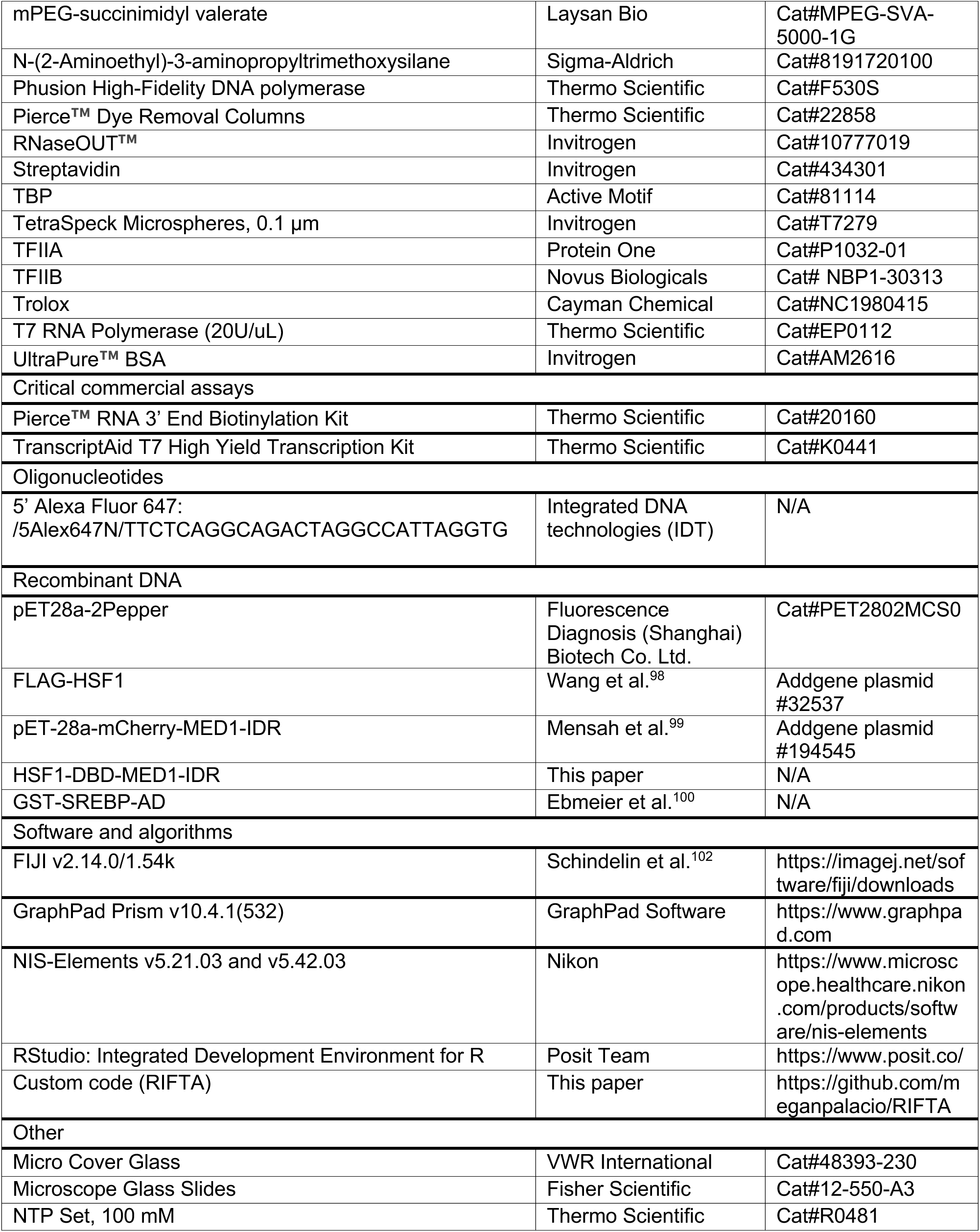

### METHOD DETAILS

#### T7 promoter DNA template

A plasmid containing an 8x-repeat of the Pepper aptamer downstream of a T7 promoter was purchased (Fluorescence Diagnosis (Shanghai) Biotech Co. Ltd. #PET2802MCS0). The T7 promoter with the Pepper array sequence was amplified by PCR (forward primer: AGGATC GAGATC TCGATC CCGC; reverse primer ATCCGG ATATAG TTCCTC CTTTCA GC) by Phusion polymerase (Thermo-Scientific #F530S). The resulting PCR product was then purified using the E.Z.N.A. Cycle Pure Kit (Omega BioTek #6492-01). The DNA template was then ethanol precipitated, washed, and resuspended to 100 nM in milliQ water, and stored frozen in single-use aliquots.

#### T7 RNAP *in vitro* transcription

The T7 promoter DNA template was mixed with 100µM HBC508 (Fluorescence Diagnosis (Shanghai) Biotech Co. Ltd. #H15081) and T7 RNAP (Thermo Scientific #EP0112) in transcription buffer (30mM Tris HCl pH 7.9, 5mM NaCl, 1mM KCl, 10mM DTT, 50µg/mL BSA, 1.67% DMSO, 0.005% Triton-X 100, 2% PEG8000). All reactions were performed in a 384 well plate (Nunc #P6491-1CS) and incubated on ice for 15 minutes. Transcription was initiated with the addition of rNTPs (2.5 mM; Thermo Scientific #R0481) and monitored at 37°C using a BioTek Synergy H1 Multimode Reader with 488nm excitation and a 15 second interval. For control RNA experiments, a DNA template had the T7 promoter followed by either a 2x- or 8x-repeat of the Pepper array was *in vitro* transcribed using the TranscriptAid T7 High Yield Transcription Kit (Thermo Scientific #K0441).

#### HSPA1B promoter DNA template

The DNA sequence of the 2x-repeat Pepper aptamer was inserted into a plasmid containing the native human HSPA1B promoter that was previously amplified from genomic DNA (HeLa) as described.^46^ The HSPA1B promoter corresponded to −501 to +100 base pairs relative to the transcription start site. The 2x-repeat Pepper DNA sequence was inserted at +101. The template was then PCR amplified using a biotinylated primer set (forward primer: /5Biosg/TTCTCA GGCAGA CTAGGC CATTAG GTG; reverse primer: GCGCGC CATTGG GATGAT) by Phusion polymerase (Thermo Scientific #F530S). For fluorescent DNA templates, a primer set of a biotinylated primer (forward primer: /5Biosg/TTCTCA GGCAGA CTAGGC CATTAG GTG) and a Cy5 primer (reverse primer: /5Cy5/GCGCGCCATTGGGATGAT) was used for amplification. The resulting PCR product was then purified using the E.Z.N.A. Cycle Pure Kit (Omega BioTek #6492-01). The DNA template was ethanol precipitated, washed, and resuspended to 100 nM in milliQ water, and stored frozen in single-use aliquots. The 19X HSF1 binding site template was generated by inserting 17 additional binding sites upstream of the −109 native HSF1 site, PCR amplified, and purified in the same manner.

#### Flow chamber functionalization

The cleaning and assembly of the flow chambers and surface functionalization were completed as previously described.^101^ Briefly, slides and coverslips were sequentially cleaned by water bath sonication using 1% alconox, followed by a 50:50 methanol-ethanol solution, then 1M potassium hydroxide, and finally 100% methanol. Slides and coverslips were then coated with 2% aminosilane before pegylation with 0.38% biotin-PEG-succinimidyl carbonate (w/v) and 20% mPEG-succinimidyl valerate (w/v) in 0.1M sodium bicarbonate. Next, slides and coverslips are assembled into flow chambers and stored at room temperature in a light-blocking container with desiccant.

### Real-time *in vitro* fluorescent transcription (RIFT)

#### Flow chamber preparation (immediately before RIFT)

Prior to RIFT, the flow chamber was washed twice with milliQ water, followed by filtered DB/RM buffer (10 mM Tris-HCl pH 7.9, 50 mM KCl, 4 mM MgCl2, 10 mM HEPES pH 7.9, 10% glycerol, 1 mM DTT, 0.05 mg/mL BSA). Streptavidin (0.2 mg/mL) was bound to the biotinylated slide surface in DB/RM buffer with 0.08 mg/mL BSA. After a 5-minute incubation, the slide was washed again with DB/RM. At this point, the flow chamber was placed into the microscope slide holder stage which was encased in an environmental chamber set to 30°C. The field-of-view (FOV) was photobleached for 1 minute to remove background noise.

#### Reconstituted in vitro transcription for “standard condition” experiments

For standard RIFT conditions, the HSPA1B promoter template (final concentration of 0.06nM) was incubated with HSF1 (13 nM, unless otherwise stated) in DB buffer to make the ‘template mix’ (20 mM Tris-HCl pH 7.9, 100 mM KCl, 20% glycerol, 1 mM DTT, 0.1 mg/mL BSA) at 30°C for 5 minutes. Next, PIC components (TFIIA, TFIIB, TFIID (or TBP), TFIIE, TFIIF, TFIIH, RNAPII, and Mediator) were added with HBC525 (final concentration of 1 µm; Fluorescence Diagnosis (Shanghai) Biotech Co. Ltd. #H15251) to the template mix in order to make the ‘PIC mix’. The concentration of PIC factors used in the assay were empirically determined but were approximately 10-40nm, similar to prior ensemble assays.^45,46^ The PIC mix was incubated at 30°C for 5 minutes, then flowed across the prepared slide, and incubated for another 5 minutes for template immobilization. A short time series (50 frames) was collected at this point to obtain images that would serve to evaluate noise/background. These images was collected for all replicates to correct for experiment-to-experiment variability. Transcription was initiated by adding A/C/G/UTP (final concentration of 2.5 mM) in imaging buffer (3 mM Trolox, 1 mg/mL glucose oxidase, 0.08 mg/mL catalase, 0.8% D-glucose, 1.7 mM RNaseOUT in DB/RM buffer; prepared as described)^101^ onto the flow chamber.

#### Imaging transcription in real time

Data collection (3-minute continuous imaging with no interval delay and 200ms exposures) started prior to NTP addition so that t=0 was captured. For two-color imaging experiments where both RNA and TF where imaged simultaneously, the exposure was 400ms due to alternating laser excitation. However, there were slight delays (∼3-6 sec) after NTP addition caused by the micro-adjustments of z-height required to achieve proper focus. All experiments were completed with two biological replicates (n=2), except for “Full PIC” results (n=6).

#### Pre-bound HSF1 versus NTP TF “stimulus response” experiments

The RIFT protocol described above was for standard RIFT in which pre-bound HSF1 was incubated with the DNA template prior to PIC assembly and NTP addition. To mimic a stimulus response condition, the TF (e.g. HSF1, GST-SREBP-AD, or DBD-MED1-IDR) was added with NTPs. In these cases, the RIFT assay followed the same protocol as standard conditions except that i) the HSPA1B promoter template (final concentration of 0.06nM in DB buffer) was incubated without TF at 30°C for 5 minutes prior to the assembly of PICs on the template and ii) transcription was initiated by adding a mix of rNTPs (final concentration 2.5mM) and TF (13nM, unless otherwise stated) in imaging buffer onto the flow chamber.

#### Imaging controls

Sets of control experiments were completed to image fluorescent microspheres (Invitrogen #T7279) for microscope calibration. Experiments with fluorescent HSPA1B promoter template (0.06 nM; labeled by Cy5 as described above) were completed to confirm template immobilization and spacing of templates on the slide surface. Purified fluorescent RNA (0.1µg; *in vitro* transcribed by T7 RNAP as described above) containing the 2x-Pepper aptamer was attached to a biotin moiety by RNA 3’-end labeling (Thermo Scientific #20160). smTIRF imaging of the purified RNA with HBC525 ligand (1 µM) validated the stability of the fluorophore:aptamer complex, and standardized fluorescence intensity of single RNAs produced during RIFT experiments.

### Single-molecule data collection and analysis

#### smTIRF microscopy

Flow chambers were imaged using an objective-based TIRF microscope (Nikon Ti2-E) equipped with a 100x 1.49NA Apo TIRF oil objective, Piezo Z stage, Oko Labs full enclosure environmental chamber, automated shutter system, and CCD camera. Manual TIRF alignment was performed prior to every experiment. Samples were excited with 488nm, 561nm, or 647nm lasers or a combination of compatible lines depending on the experiment. Emission was captured by the Andor Ixon 897 EMCCD. Data were collected through the NIS elements software (version 5.21.03) at 200ms exposure time with no interval delay for the duration of the movie.

Early on in development of RIFT, experiments were conducted using a different TIRF microscope (Nikon TE-2000 U), which featured a 1.49NA oil immersion objective, a Piezo nano-positioning stage, and two CCD cameras. Emission from 532nm excitation was captured with an Evolve Photometric CCD, while emission from 635nm excitation was captured with a Cascade II Photometric CCD. However, this setup posed limitations on imaging near t=0, prompting a transition to a more advanced TIRF microscope (Nikon Ti2-E; described above). All data presented were captured using the advanced Nikon Ti2-E. The sole exception was the minimal system experiment (Figure 2A), which was conducted on the Nikon TE-2000 U.

#### smTIRF microscopy analysis of RIFT data

Movies were analyzed in FIJI (ImageJ2 version 2.14.0/1.54k) to perform spot intensity analysis for single-molecule detection and particle intensity quantification.^102^ A low noise tolerance provided detection of all particles, including background, and their respective intensity values over time. The data were analyzed using a custom script (RIFTA) developed in R studio to determine ‘real’ transcripts from noise with high confidence (> 99^th^ percentile).^103^ The ‘PIC’ image (flow chamber with PICs and HBC ligand; no NTPs) collected for all replicates provided intensity traces of background noise or inherent fluorescence of unbound HBC ligand. The 99^th^ percentile of all intensity values for the background foci observed determined the signal filter (Figure S1F; green dashed line). This permitted the rigorous removal of intensity traces that originated from background particles that were not true RNA molecules, and provided confidence that foci that passed the filter cutoff were real transcripts. In the movies where transcription was initiated, a transcript was called when it passed the signal filter (e.g. Figure 1C). In addition, to be called a re-initiated transcript from the same promoter, the intensity values had to return to noise levels after the first transcript (e.g. Figure S1F; orange area) and subsequently surpass the filter level again. This same data processing protocol was applied for every experiment to account for variance. Moreover, the RIFTA script reported amounts and rates of first round PIC activation and re-initiation.

#### smTIRF microscopy analysis of fluorescent TFs

Movies were analyzed in FIJI (ImageJ2 version 2.14.0/1.54k) to perform spot intensity analysis for single-molecule detection. Additionally, TF foci were distinguished from background by setting a threshold, and measurements of foci size and intensity were taken. Foci consisting of multiple molecules (clusters) versus single molecules were sorted based upon fluorescence intensity. Additionally, TF dwell times were calculated by measuring the duration a TF remained at the same location.

### Confocal microscopy

Confocal microscopy images were collected on a spinning disc confocal (Nikon TiE) equipped with a 100x 1.4NA oil objective, MCL Piezo Z stage, environmental control (Oko Labs enclosure), and Andor IXon 888 Ultra camera. Samples were excited with 488nm, 561nm, or 640nm lasers or a combination of compatible lines depending on the experiment. Movies were captured through the NIS elements software (version 5.42.03) with no interval delay for the duration. Data were then analyzed in FIJI (ImageJ2 version 2.14.0/1.54k) to minimize background, detect all particles, and quantify particle size and intensity across all movies. mCherry-MED1-IDR (200nM), DBD-MED1-IDR (100nM), and HSF1 (100nM) were imaged in DB/RM buffer at 30°C.

### Purification of human PIC factors

TFIID, TFIIE, TFIIF, TFIIH, Mediator, and RNAPII were purified as described.^46^ The remaining GTFs were purchased: TFIIA (Protein One #P1032-01), TBP (Active Motif #81114), TFIIB (Novus Biologicals #NBP1-30313).

### Purification of HSF1, GST-SREBP-AD, MED1-IDR, and DBD-MED1-IDR chimera

mCherry-MED1-IDR and mCherry-DBD-MED1-IDR were expressed in *E.coli* LOBSTR (low background strain; Kerafast #EC1002) cells. Cells were grown at 37°C to an OD ∼ 0.7, induced with 1mM IPTG (American Bio #AB00841), and continued growth for 4 hours at 37°C. Both proteins had a 6xHis tag allowing for purification by nickel affinity chromatography using a HisTrap HP column (Cytiva # 17524701). Peak fractions were then purified further with a Superdex 200 Increase 10/300 GL (Cytiva # 28990944) size exclusion column. GST-SREBP-AD was expressed in *E.coli* and cells were grown at 37°C to an OD ∼0.7, induced with 1mM IPTG, and continued growth for 3 hours at 30°C. GST-SREBP-AD was isolated from lysate by binding to glutathione-sepharose beads (Cytiva #17075601) and eluted with buffer containing 30 mM glutathione (Sigma-Aldrich #G4251). HSF1 was expressed in Expi293F (Gibco #A14527) cells that underwent 0.5-hour heat shock (42°C) after 48 hours transfection. The HSF1 construct had a FLAG tag that allowed purification using FLAG M2 affinity resin (Sigma-Aldrich #A2220-5ML) from whole cell extracts. HSF1 was eluted with 5mg/mL of FLAG peptide (Sigma-Aldrich #F4799-25MG) in 0.15M KCl HEGN.

### Fluorescent labeling of TFs HSF1 and GST-SREBP-AD

For microscopy experiments visualizing TFs, HSF1 and GST-SREBP-AD were fluorescently labeled with Alexa Fluor™ 647 (Invitrogen #A20347). Alexa Fluor labeling was completed by incubating samples with a 10X molar excess of TCEP (tris-(2-carboxyethyl)phosphine; Invitrogen #T2556) and 2X molar excess of Alexa Fluor 647 for 2 hours at 4°C while nutating and protecting from light. Excess dye was removed using Pierce dye removal columns (Thermo Scientific #22858). Labeling efficiency was assessed using gel electrophoresis, with BSA and Alexa Fluor 647 oligo standards serving to determine the protein concentration of the sample and the extent of fluorescence incorporation.

## QUANTIFICATION AND STATISTICAL ANALYSIS

### Probability of HBC:aptamer dissociation

The affinity of HBC525 for the Peppers aptamer is Kd = 3.8 nM; HBC ligands have slow dissociation rates (0.0023 s^-1^) and high photostability (>3600 s).^44^ With the assumption that one imaging FOV (field of view; ca. 800 templates) produced 300 transcripts and that ligand-aptamer binding is in equilibrium, we estimate that the probability of a single aptamer not being bound by HBC525 is 0.38% and the probability of both aptamers in the 2x array remaining unbound is 0.0014%. Thus, the probability that an RNA transcript would remain ‘dark’ and undetected was low.

### Calculation of burst duration

Burst durations from RIFT data were calculated by determining the slope of total transcription over time, tiling across the entire experimental timeframe (t=3min). A burst was considered to have ended when the slope dropped below 0.167 (equivalent to fewer than 1 transcript/6 seconds). The corresponding time point at which this occurred was recorded as the burst duration. Data confirmed that transcription past the burst duration was minimal. Some experiments were unable to provide a burst duration value because the burst continued through the experiment, indicating that the burst duration in these cases was at least 3min.

### Statistical analysis

Statistical comparison of RIFT data was performed using a Welch’s t-test, to determine p-values while accounting for variance in sample sizes. P-values are reported on plots and the “ns” label was used for any p-value ≥ 0.2. Outliers were determined by robust regression and outlier removal (ROUT method) with a strict Q = 0.1%.^104^ Statistical analyses and plot generation was completed using GraphPad Prism 10 for macOS version 10.4.1(532).

