## Supplemental Figures for "Real-time visualization of reconstituted transcription reveals RNA polymerase II activation mechanisms at single promoters"

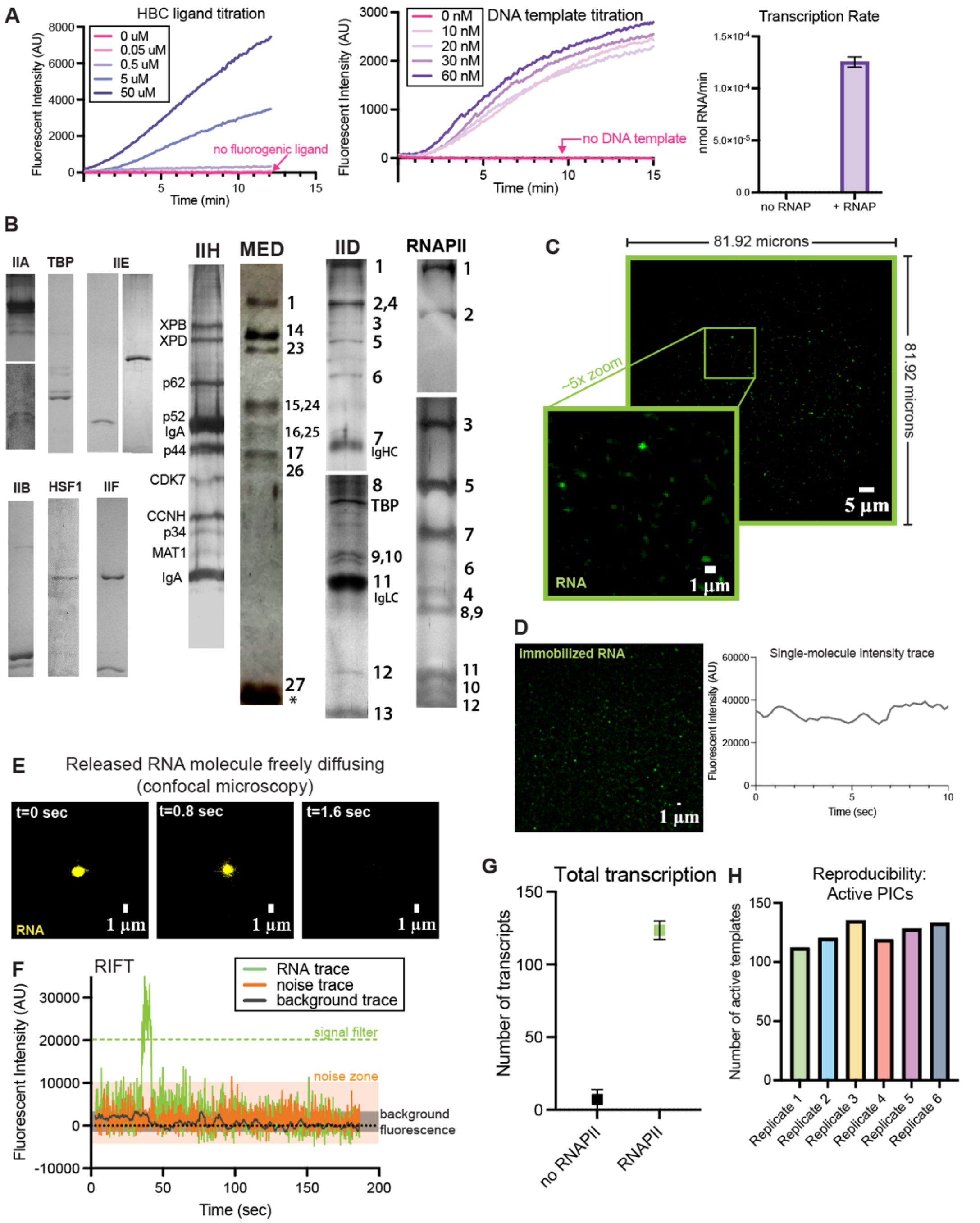


**Figure S1. Additional details and controls for RIFT (related to Figure 1)**

(A) T7 RNAP transcription on DNA template containing 8x-Pepper aptamer with HBC508. Bulk fluorescence measured by plate reader confirmed signal was dependent on fluorogenic ligand (left), DNA (middle), and RNAP (right) (n=3 biological replicates).

(B) Purified human PIC factors used for RIFT. Some samples (TFIIA, IID, RNAPII) were run on two gels with different acrylamide percentages to resolve small proteins. Bands for TFIIA: 56 & 13kDa; TFIIE: 34 & 56kDa; TFIIF: 74 & 30kDa. For Mediator, TFIID, and RNAPII, subunits are abbreviated to save space (e.g. MED1, MED14, MED23; TAF1, TAF2, TAF4; RPB1, RPB2, and so on). For Mediator, smaller proteins were not resolved on the 7% gel (asterisk: GST-SREBP-AD). Gels for TFIIA, TFIID, IIE56, TFIIH, Mediator and RNAPII are silver-stained and the others are coomassie-stained. Note silver staining is not quantitative; band intensities will depend on the number of Cys residues in the protein.

(C) Field-of-view (FOV) and imaging dimensions of RIFT experiments. Imaging over an area captures hundreds of single molecules which permits evaluation of population effects in addition to single molecules.
(D) smTIRF image of immobilized biotinylated 2x-Pepper RNA (+HBC525 ligand) and representative single RNA intensity trace (n=2 biological replicates).
(E) Representative example of freely diffusing "runoff" RNA (+HBC525 ligand) produced from RIFT; imaged by confocal microscopy.
(F) Example RIFT intensity traces of nascent RNA (green line), noise (orange line; assumed to be free HBC525 ligand), and background (black line). Any intensity trace that did not surpass the filter was discarded.
(G) Total transcription as a function of RNAPII (n=2 biological replicates).
(H) Reproducibility of RIFT across multiple independent experiments (n=6).

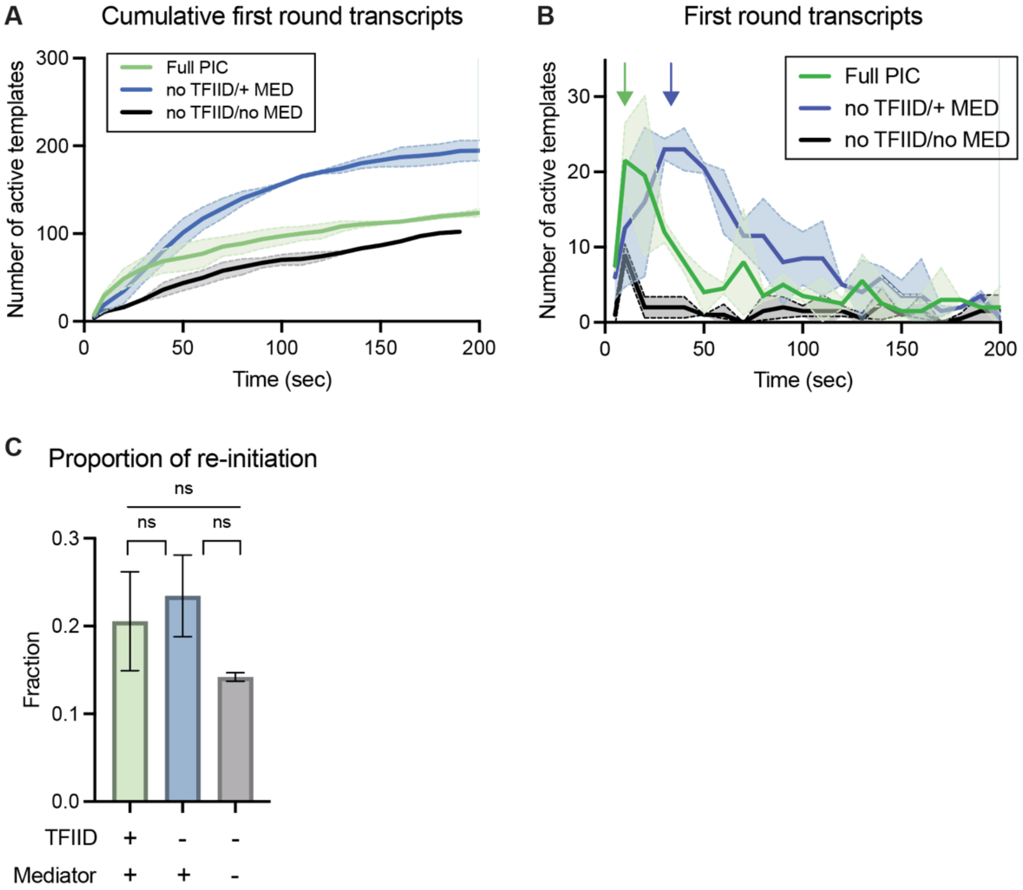


**Figure S2. TFIID accelerates RNAPII activation (related to Figure 2)**
(A) Cumulative first round transcripts. Note that "no TFIID" experiments include TBP. Line depicts mean values and shading represents SEM.
(B) Number of first round transcripts over time. Green arrow highlights the faster activation with the full PIC compared with PICs lacking TFIID (blue arrow). Note that "no TFIID" experiments include TBP. Line depicts mean values and shading represents SEM.
(C) Fraction of active promoters that undergo re-initiation. Note that "no TFIID" experiments include TBP. Bars represent mean ± SEM.
Data panels A-C represent n=2 biological replicates.

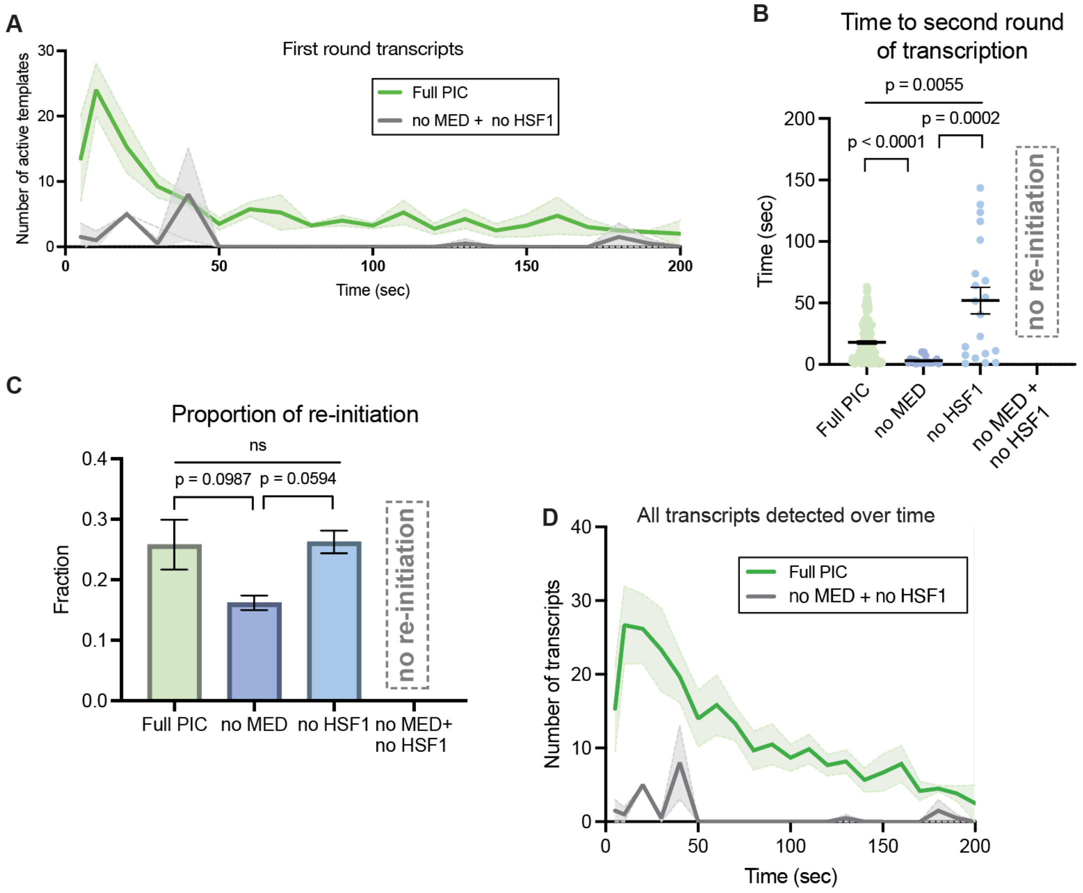


**Figure S3. HSF1 and Mediator cooperatively activate RNAPII (related to Figure 3)**

(A) Number of first round transcripts over time ±Mediator and HSF1. Line depicts mean values and shading represents SEM.

(B) Time to re-initiation (second round) based upon PIC composition. Black bars represent mean ± SEM.

(C) Fraction of active promoters that undergo re-initiation based on Mediator, HSF1 or both factors. Bars represent mean ± SEM.

(D) Number of transcripts over time ±Mediator and HSF1. Line depicts mean values and shading represents SEM.

All panels shown (A-D) represent data from 2 biological replicates, except n=6 for Full PIC experiments.

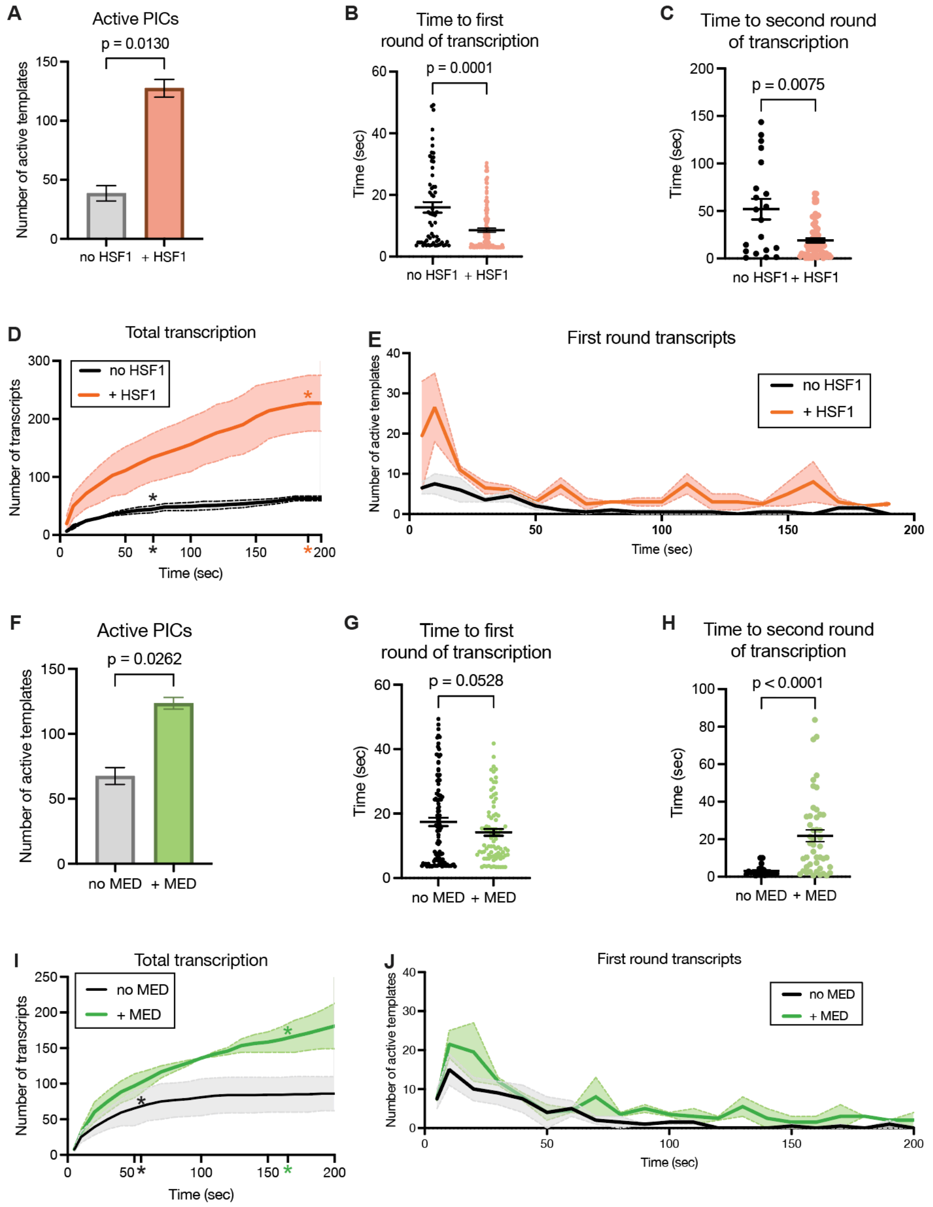


**Figure S4. HSF1 and Mediator independently increase RNAPII transcription (related to Figure 3)**

(A) Number of active PICs ±HSF1. Bars represent mean ± SEM.
(B) Time to first round transcription as a function of HSF1. Black bars represent mean ± SEM.
(C) Time to re-initiation (second round) ±HSF1. Black bars represent mean ± SEM.
(D) Total transcription over time as a function of HSF1. Asterisks signify burst duration, line depicts mean values, and shading represents SEM.
(E) Number of first round transcripts plotted over time, as a function HSF1. Line depicts mean values and shading represents SEM.
(F) Number of active PICs ±Mediator. Bars represent mean ± SEM.
(G) Time to first round transcription as a function of Mediator. Black bars represent mean ± SEM.
(H) Time to re-initiation (second round) ±Mediator. Black bars represent mean ± SEM.
(I) Total transcription over time as a function of Mediator. Asterisks signify burst duration, line depicts mean values, and shading represents SEM.
(J) Number of first round transcripts plotted over time, as a function of Mediator. Line depicts mean values and shading represents SEM.
Data panels A-J represent n=2 biological replicates.

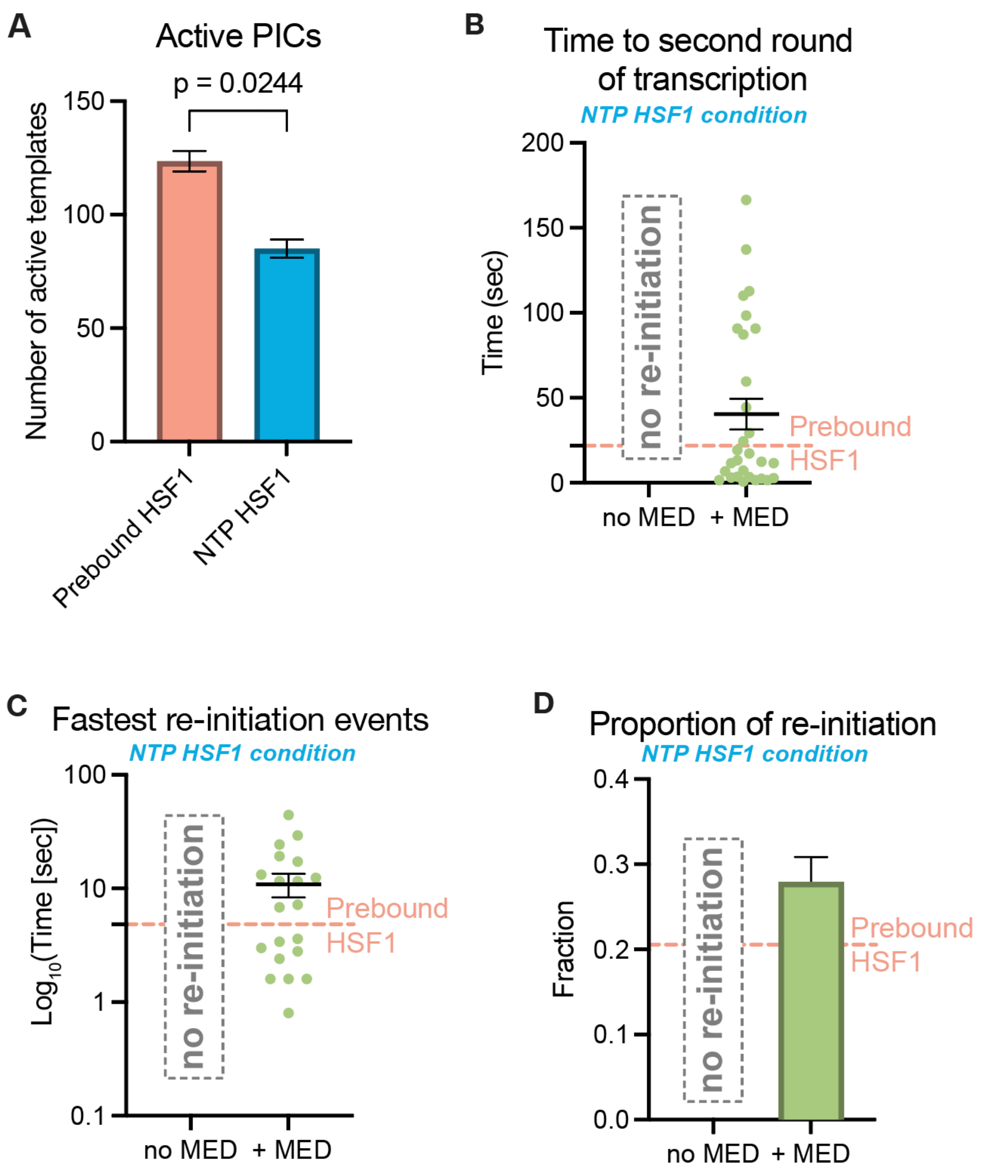


**Figure S5. Additional data for HSF1- and Mediator-dependent RNAPII activation under stimulus response experimental conditions (related to Figure 4)**

(A) Number of active PICs from pre-bound HSF1 or NTP HSF1 conditions. Bars represent mean ± SEM.

(B) Time to re-initiation (second round) ±Mediator in NTP HSF1 condition. Re-initiation did not occur in the absence of Mediator for NTP HSF1 condition. Dashed orange line conveys pre-bound HSF1 results for comparison. Black bars represent mean ± SEM.

(C) Time to re-initiation for the fastest events (n=20) ±Mediator under the NTP HSF1 condition. Dashed orange line conveys pre-bound HSF1 results for comparison. Black bars represent mean ± SEM.

(D) Fraction of active promoters that undergo re-initiation ±Mediator for the NTP HSF1 condition. Bars represent mean ± SEM.

Data panels A-D represent n=2 biological replicates.

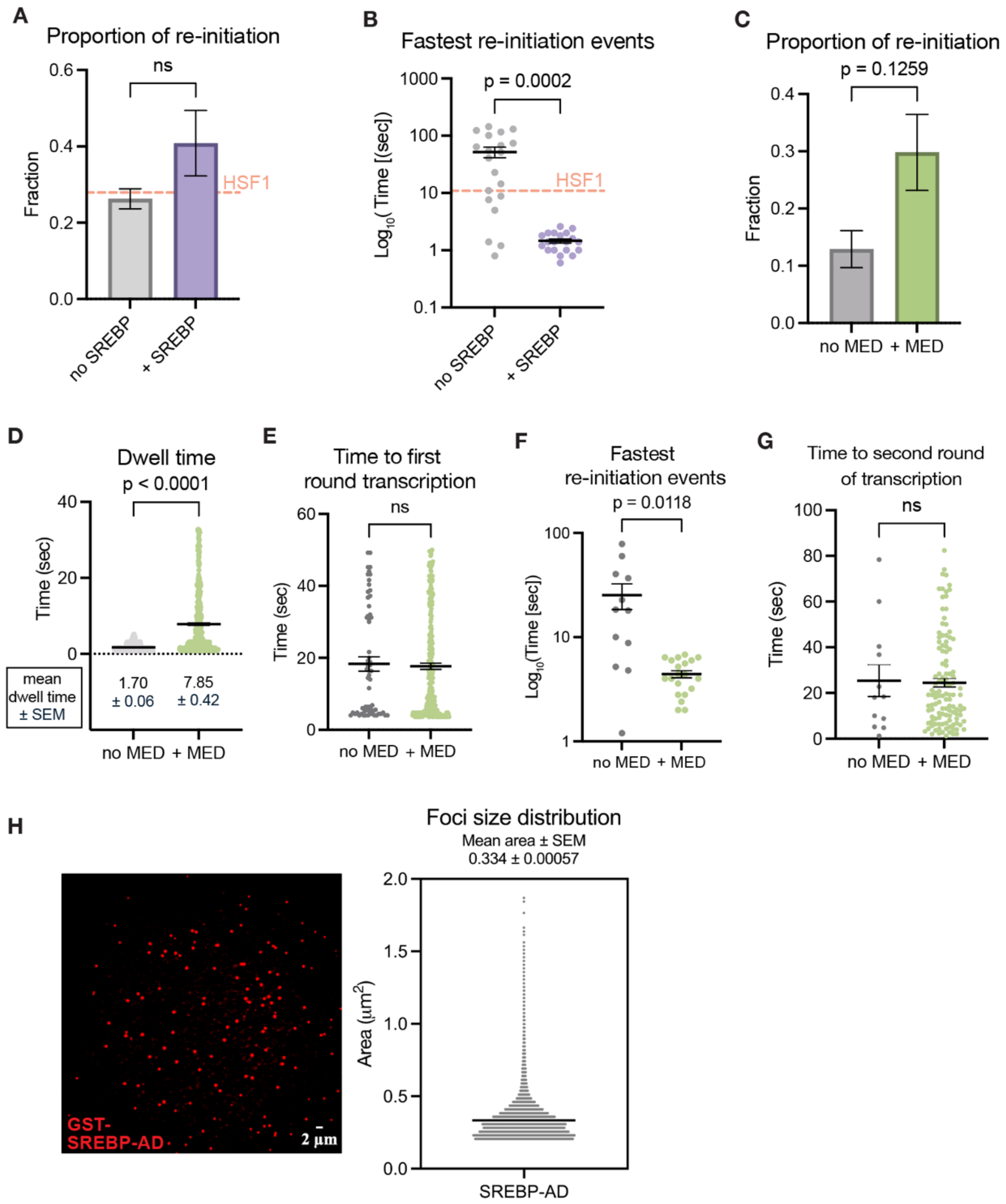


**Figure S6. Mediator enables rapid TF-dependent RNAPII activation without TF-DNA binding or condensate formation (related to Figure 5)**

(A) Fraction of active promoters that undergo re-initiation ±GST-SREBP-AD, which was added with NTPs in these assays. Dashed orange line shows results from NTP HSF1 condition for comparison. Bars represent mean ± SEM.

(B) Time to re-initiation of the fastest events (n=20) ±GST-SREBP-AD. Dashed orange line conveys NTP HSF1 results for comparison. Black bars represent mean ± SEM.

(C) Fraction of active promoters that undergo re-initiation ±Mediator in NTP GST-SREBP-AD condition. Bars represent mean ± SEM.

(D) Dwell time of fluorescently labeled (AlexaFluor647) GST-SREBP-AD as a function of Mediator. Black bars represent mean ± SEM, and mean values ± SEM are shown.

(E) Time to first round transcription as a function of Mediator with the NTP GST-SREBP-AD condition. Black bars represent mean ± SEM.

(F) Fastest re-initiation events (n=20) ±Mediator with the NTP GST-SREBP-AD condition. Black bars represent mean ±SEM.

(G) Time to re-initiation (second round) ±Mediator with the NTP GST-SREBP-AD condition. Black bars represent mean ±SEM.
(H) Representative image of fluorescently labeled (AlexaFluor647) GST-SREBP-AD during RIFT. The corresponding foci size distribution of GST-SREBP-AD also shown. Black bars represent mean ± SEM.
Data panels A-H represent n=2 biological replicates.

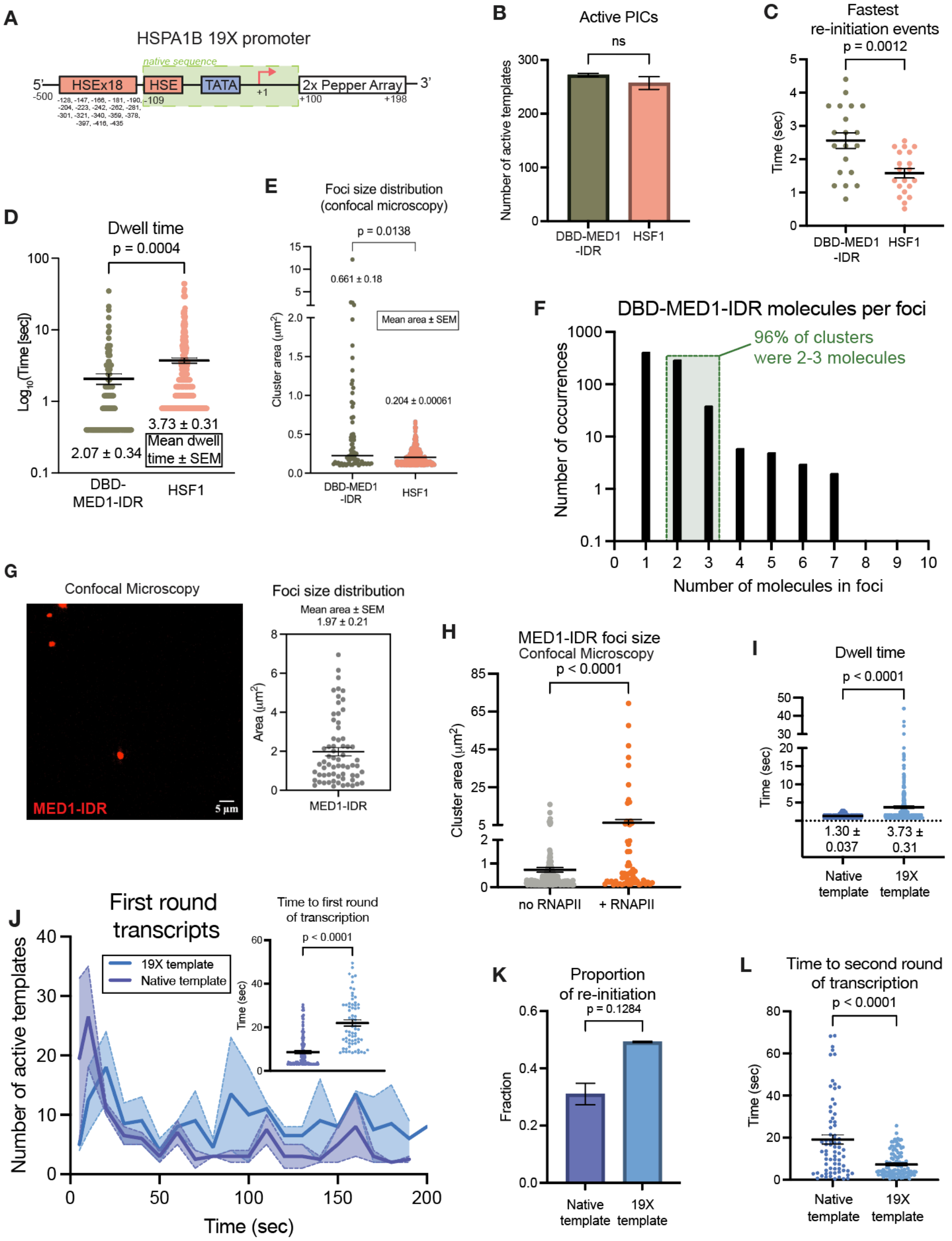


**Figure S7. MED1-IDR can replace HSF1 to activate RNAPII; promoter occupancy drives RNAPII burst size and burst duration (related to Figure 6 and 7)**

(A) Schematic of 19X HSPA1B template with conserved native sequence highlighted in green. Locations of inserted HSF1 binding sites are shown.

(B) Number of active PICs as a function of DBD-MED1-IDR or HSF1. Bars represent mean ±SEM.

(C) Fastest re-initiation events (n=20) for DBD-MED1-IDR or HSF1. Black bars represent mean ±SEM.

(D) Dwell time of fluorescently labeled (AlexaFluor647) DBD-MED1-IDR or HSF1. Black bars represent mean ±SEM, and mean values ±SEM are shown.

(E) Confocal microscopy of fluorescent DBD-MED1-IDR (100nM) or HSF1 (100nM) to measure foci size distribution. Black bars represent mean ±SEM, and mean values ±SEM are shown.

(F) Distribution frequency of the number of DBD-MED1-IDR molecules per spot. The majority of DBD-MED1-IDR foci were single molecules and nearly all clusters consisted of 2-3 molecules (green box). Note y-axis is Log10 scale.

(G) Example of MED1-IDR (200nM) imaged by confocal microscopy. At right is size distribution of foci. Black bar represents mean ±SEM and the mean value is shown.
(H) MED1-IDR foci size ±RNAPII imaged by confocal microscopy. The increased spot size +RNAPII suggests RNAPII partitioning with MED1-IDR and is consistent with condensate behavior. Black bars represent mean ± SEM.
(I) Fluorescently labeled HSF1 (100 nM; AlexaFluor647) dwell time shown on the native or 19X HSPA1B template. Black bars represent mean ±SEM and mean values are shown below.
(J) Number of first round transcripts plotted over time at the native or 19X HSPA1B promoter. Line depicts mean values and shading represents SEM. Inset panel displays time to first round transcription of native or 19X HSPA1B promoter, using the same color scheme. Black bars represent mean ±SEM.
(K) Fraction of active promoters that undergo re-initiation from the native or 19X HSPA1B promoter. Bars represent mean ± SEM.
(L) Time to re-initiation (second round) from the native or 19X HSPA1B promoter. Black bars represent mean ±SEM.
Data panels in B-L represent n=2 biological replicates.
